# GPR17 modulates oligodendrocyte precursor cell maturation during development but has limited impact on myelin regeneration following demyelinating insults

**DOI:** 10.1101/2025.07.08.659146

**Authors:** Kristin Puorro-Radzwill, Petti Pang, Stefanie Giera, Guoqing Sheng, Jonathan Farley, Carrie Garron, Sanjay Danthi, Joseph Gans, Dinesh Kumar, Rajiv Agrawal, Dinesh Bangari, Susan Ryan, Nilesh Pande, Tarek A. Samad, Sabine Gratzer, Carlos Pedraza

## Abstract

Pharmacological enhancement of myelin regeneration is broadly recognized as the next frontier in therapeutic approaches for demyelinating diseases of the CNS such as multiple sclerosis. However, although several molecular targets for remyelination have been tested preclinically and in clinical trials, an efficacious and safe myelin repair treatment is yet to be developed. One promising molecular target to enhance myelin repair is the G protein-coupled receptor (GPCR) GPR17, which has been proposed to play a central role in the transition from early oligodendrocyte progenitor cells (OPC) into pre-myelinating oligodendrocytes. These findings are largely supported by studies using transgenic mice where GPR17 deletion results in developmental hypermyelination. Additionally, pharmacological modulation of GPR17 activity has been reported to enhance oligodendrocyte precursor cell (OPC) maturation and myelination. In our studies aimed to characterize and pharmacologically validate GPR17 as a viable target for drug development, we established by means of transcriptional profiling of GPR17 knockout versus wild type OPCs, that absence of this GPCR results in a gene signature revealing minor changes in myelin protein gene expression. Furthermore, blocking GPR17 receptor activity in OPC cultures using selective and potent antagonists or inverse agonists, results in limited enhancement of maturation and myelination *in vitro*. Importantly, remyelination in both the cuprizone and lysolecithin-induced demyelination models was not enhanced in the absence of GPR17. Our data demonstrate that GPR17 plays a minor role in OPC differentiation during development, and pharmacological modulation of its activity has a marginal effect on oligodendrocyte precursor maturation and myelin regeneration after injury.

## Introduction

Myelin repair in early stages of multiple sclerosis (MS) occurs spontaneously as lesions, caused by immune cell attacks, resolve. This natural regeneration mechanism slows down to a halt in advanced stages of the disease, eventually resulting in axonal degeneration, brain atrophy, accumulation of lesions and neurological disabilities (Fancy et al., 2010; Motavaf, Sadeghizadeh, & Javan, 2017). Available immunotherapies to treat MS, while effectively decreasing relapse rates and slowing down disease progression, fail to significantly prevent axonal injury and degeneration and to improve myelin repair (Rommer et al., 2019). Importantly, remyelination can be enhanced pharmacologically, as demonstrated in preclinical models and in clinical trials, by stimulating resident oligodendrocyte progenitors (OPCs) to mature into myelinating oligodendrocytes (Green et al., 2017; Schwartzback et al., 2017; Zhang et al., 2012). Therefore, novel MS therapies currently in research and development are focused on enhancing remyelination and neuronal protection as adjunctive treatments to regulators of the autoimmune response (Coclitu, Constantinescu, & Tanasescu, 2016; Kolahdouzan, Futhey, Kieran, & Healy, 2019).

Several signaling pathways and molecular targets have been proposed to modulate OPC maturation as potential therapies for remyelination (Hooijmans et al., 2019; Olsen, & Akirav, 2015; Stangel, Kuhlmann, Matthews, & Kilpatrick, 2017). Among them, GPR17, a G protein-coupled seven transmembrane domain receptor, has been reported to be a central modulator of OPC maturation from early bipolar OPC stages into multi-branched, pre-myelinating oligodendrocytes (Chen et al., 2009; Dziedzic, Miller, Saluk-Bijak, & Bijak, 2020; Hennen et al., 2013; Ou et al., 2016). GPR17 expression profile correlates with a putative role in OPC maturation as its expression is restricted to early progenitors, peaking when the cells transition from bipolar morphology to multi-branched cells with membranous structures (Fumagalli et al., 2011; Hennen et al., 2013). Additionally, genetic deletion of GPR17 in the mouse was associated with early onset of developmental myelination while overexpression of GPR17 resulted in hypomyelination (Chen et al., 2009; Ou et al., 2016). Interestingly, while overexpression of the receptor under the myelin protein CNP1 promoter was postnatally lethal, the GPR17 knockout mice did not exhibit overt phenotypes and the global early increase in myelin content reverted to normal levels before adulthood when compared to wild type brains. This expression pattern indicated a predominant role of the receptor during a short time window of OPC maturation and effectively erected modulation of GPR17 activity as a potential therapy to enhance myelin repair. Yet, conclusive evidence of pharmacological modulation of GPR17 activity, with selective and potent ligands, resulting in enhanced OPC maturation and remyelination after injury is still controversial. (Parravicini et al., 2020)

In order to determine the druggability of GPR17 as a viable remyelination therapy, we have characterized genetic and pharmacological modulation of the receptor *in vitro* and *in vivo* to assess its role in regulating OPC maturation and myelination during development and following demyelinating injury in adult mice. Our data demonstrate that lack of GPR17 expression in cultured OPCs results in a distinct transcriptional signature with enhanced expression of genes involved in cellular processes such as axonal guidance, gap junction and cholesterol biosynthesis pathways. However, limited association with myelin formation gene signatures was detected in the absence of GPR17. We have also evaluated the impact of modulating the receptor’s activity *in vitro* with highly selective and potent antagonists, agonists and inverse agonists by assessing the level and speed of OPC maturation. *In vivo*, remyelination in two different animal models of demyelination was not significantly enhanced in GPR17 deficient mice. Our findings support a transitional, minor role for GPR17 in OPC biology during developmental myelination with no measurable influence on myelin repair following demyelinating injury in the adult mouse.

## Methods

### Animals

Experimental protocols were approved by Sanofi’s institutional animal care and use committee, and studies were conducted in Sanofi’s association for assessment and accreditation of laboratory animal care accredited facility.

### Primary rat oligodendrocyte progenitor cells

Whole brains from postnatal day 1-3 Sprague-Dawley rats were dissociated using the Miltenyi neural tissue dissociation kit according to the manufacturer’s instructions (Miltenyi, Bergisch Gladbach, Germany). Oligodendrocyte precursor cells (OPCs) were purified using magnetic activated cell sorting (Miltenyi MACS) technology with anti-A2B5 microbeads (Miltenyi Biotec, Cat# 130-093-388, RRID:AB_2833097) according to the manufacturer’s protocol and were cultured on Cell Start (Life Technologies, Carlsbad, USA) coated tissue culture flasks in DMEM/F12 media supplemented with 1X B27 minus Vitamin A, 10 ng/mL each of recombinant human basic FGF and PDGF-AA (Life Technologies) 100 units/mL of penicillin and 100 µg/mL of streptomycin (proliferation media). Cells were maintained at 37⁰C in a humidified incubator enclosing an atmosphere of 95% air/5% CO_2_ for 3-5 days.

### Rat OPC cell maturation assay and compound treatment

Purified rat OPCs were plated in OPC differentiation media (DMEM/F12, 1X B27 minus Vitamin A, 2 ng/ml PDGF-AA, 2 ng/ml basic FGF, and 100 units/mL of penicillin and 100 µg/mL of streptomycin) at 10^4^ cells/well onto poly-D-lysine (PDL) (Sigma-Aldrich, St. Louis, USA) coated 96 well plates. 24 hours later the cells were treated with compounds supplied in 10mM DMSO stock solutions and diluted into indicated final concentrations in differentiation media. The final DMSO concentration across all treatments, including vehicle, was 0.1%. Cells were harvested for analysis after 3- or 6-days of treatment. For 6 days of compound exposure an intermediate treatment was performed at day 4 by removing a third of the media from the wells and adding compounds in fresh media solutions.

### Primary mouse oligodendrocyte progenitor cells

Mouse oligodendrocyte precursor cells (OPCs) were generated from postnatal day 1-3 brains from wild type (Wt, GPR17 ^+/+^), heterozygotes (Het, GPR17 ^+/-^) or homozygotes (Hom, GPR17 ^-/-^) following a protocol by O’Meara, Ryan, Colognato, & Kothary, (2011) with some modifications. Whole brains were dissected and dissociated using Miltenyi’s neural tissue dissociation kit according to the manufacturer. Following dissociation, cells were filtered through a 70μM filter and plated in DMEM+10% fetal bovine serum+100U/mL of penicillin and 100μg/mL of streptomycin (all from Life Technologies) on 100μg/mL poly-D-lysine (Sigma) coated T75 tissue culture flasks (non-vented) at 3-4 brains/flask. Mixed glial cultures were cultured for eight to ten days at 37°C, 5% CO_2_. Every 2-3 days, a 2/3 media change was performed. On day 7, the media was supplemented with human insulin (Sigma) to a final concentration of 5μg/mL. On day 8, in order to remove loosely attached, proliferating microglial cells, the flasks were shaken for 45 minutes at 50rpm on a benchtop orbital shaker set at 37°C. Media was replaced with human insulin supplemented (5μg/mL final) culture media and the flasks were returned to the incubator. After an hour, the flasks were placed again on the orbital shaker set at 220rpm overnight. The next day, dislodged OPCs were harvested by collecting the supernatant and plating for differential adhesion on to a non-tissue culture treated 15cm dish (ThermoFisher, Waltham, USA) for 30 minutes. At the 15-minute mark, the dishes were given a gentle nudge to prevent any OPCs from adhering to the plastic. Non-adherent cells were collected, counted and seeded onto Cell Start (Life Technologies) coated T75 tissue culture flasks at 5x10^6^ per flask in SATO media (Dugas, & Emery, 2013) containing DMEM with 4.5g/L glucose, 2mM glutamine, 100U/mL penicillin, 100μg/mL streptomycin, 1mM sodium pyruvate, 1X B27 without vitamin A, 20ng/mL human PDGF-AA (all from Life Technologies), 5μg/mL human insulin, 5μg/mL N-Acetyl-L-cysteine, 10ng/mL d-biotin, 100μg/mL bovine serum albumin, 100μg/mL apo-transferrin, 16μg/mL putrescine, 60ng/mL progesterone, 40ng/mL sodium selenite, 4.2μg/mL forskolin, (all from Sigma), 10ng/mL rat CNTF, 1ng/mL human NT-3 (both from Peprotech) and 1X Trace Elements B (Corning Life Sciences, Tewksbury, USA). Cells were incubated at 37°C, 10% CO_2_. A full media change was performed the next day. Every two days thereafter, half of the media was removed and replaced with fresh media. Cells were ready for plating and subsequent experiments by 3-5 days *in vitro*.

### Enzyme-linked immunosorbent assay (ELISA) analysis of myelin basic protein (MBP)

Standard sandwich ELISA was performed using the following antibodies diluted in PBS (coating antibody) or PBS containing 1% bovine serum albumin (all other antibodies). Coating antibody: monoclonal anti-MBP (1:2500, Millipore Cat# MAB382, RRID:AB_94971); detection antibody: polyclonal anti-MBP (1:2500, Abcam Cat# ab28541, RRID:AB_776581); biotinylated goat anti-rabbit IgG antibody (1:10,000, Vector Laboratories Cat# BA-1000, RRID:AB_2313606), streptavidin-biotinylated HRP complex (1:8000, GE Healthcare, Chicago, USA). Cells harvested at 3- and 6-days post treatment were lysed in triple detergent buffer (50 mM Tris-HCl, pH 8.0, 150 mM sodium chloride, 0.02% sodium azide, 0.1% sodium dodecyl sulfate, 1.0% NP-40, 0.5% sodium deoxycholate, all from Sigma) containing 1X Complete Mini, EDTA-free Protease Inhibitor Cocktail (Roche, Mannheim, Germany). Known concentrations of recombinant bovine MBP (Invitrogen) were used to generate a standard curve. Standards and cell lysates were added to 96-well Maxisorp plates (Nunc, ThermoFisher) pre-coated with coating antibody and incubated overnight at 4°C. Plates were washed three times with phosphate buffered saline containing 0.5% Tween-20 (Sigma) (PBST) using an automated microplate washer (405 TS, BioTek Instruments Inc. Winooski, USA). Plates were then incubated at room temperature with detection antibody (2 hours), biotinylated anti-rabbit IgG (1 hour), and streptavidin-biotinylated HRP complex (1 hour) with three washes with PBST between each incubation step. To induce colorimetric change, o-phenylenediamine dihydrochloride (OPD) (Sigma) was added to each plate for 30 minutes and the reaction stopped by addition of 2N sulfuric acid (RICCA). Total MBP concentration was determined by colorimetric change of plates read at 492nm using the FlexStation® 3 Multi-Mode Microplate Reader (Molecular Devices, San Jose, USA). Total protein concentration in the lysates were determined by Bicinchoninic Acid Assay (BCA) Protein Assay (Pierce, ThermoFisher) according to the manufacturer’s instructions.

### Immunocytochemistry

Following 4- or 6-days of OPC maturation, media was aspirated, and cells were fixed with 4% formaldehyde (Electron Microscopy Sciences, Hatfield, USA) for 15 minutes at room temperature. Cells were washed three times with cold PBS and permeabilized with 0.3% Triton X-100 (Sigma) in PBS for 10 minutes at room temperature. Cells were washed three times with cold PBS and blocked with 5% normal goat serum (Millipore) in PBS containing 0.1% Tween 20 (Sigma) (PBST) for 1hr at room temperature. Primary antibody was diluted in PBST and added overnight at 4°C. Cells were washed three times in PBST and secondary antibody was diluted in PBST and added for 2hr at room temperature. Cells were washed three times in cold PBST and stained with 1μg/mL Hoechst 33342 (Life Technologies) during the last wash for 5 minutes at room temperature followed by one more wash in PBST. PBST was aspirated and replaced with PBS. Primary antibodies used for staining were: anti-GPR17 (1:100, Sigma-Aldrich Cat# HPA029766, RRID:AB_10612114), anti-MBP (1:500, Bio-Rad Cat# MCA409S, RRID:AB_325004) and anti-Nkx2.2 (1:300, DSHB Cat# 74.5A5, RRID:AB_531794).

### Whole genome RNA sequencing of mouse OPCs

Mouse OPCs were plated at 250,000 cells per well of poly-D-lysine coated six well plates in duplicate. On days 0, 1 and 3, media was aspirated, and cells lysed with buffer RLT+ (Qiagen, Hilden, Germany) supplemented with 1% 2-mercaptoethanol (Sigma) according to the manufacturer. For RNA extraction, samples were first randomized then total RNA was extracted using the Qiagen RNeasy Plus kit with optional on-column DNase digest step (all one batch). RNA quantification and quality assessment were performed using a Nanodrop 8000 and a Tapestation 4200, respectively. RNA integrity number (RIN) values ranged from 7.8-10. Total RNA was diluted to 1ng in 10μl of water then using Oligo(dT) priming from Takara’s SmartSeq v4 Ultra Low Input RNA kit, enriched mRNA was converted to full length cDNA. cDNA was amplified using 11 PCR cycles then cleaned up using AMPure XP beads. The average size of the full-length cDNA ranged from 2500-2800bp after running the Bioanalyzer 2100. Sequencing libraries were prepped and uniquely indexed using Illumina’s Nextera XT kit without modification to the standard protocol. Average size of libraries ranged from 800-900bp using the Tapestation 4200. Each sample was quantified using Qubit then diluted to 4nM and pooled together. To prepare for sequencing, pooled libraries were first denatured with NaOH then diluted to 1.8pM and finally run on an Illumina NextSeq500 using the High output kit and a 2x76bp paired end run. RNA-seq data analysis was performed using RNA-seq pipeline from the Array Suite-Omicsoft Corporation, (Version 6.2, Array Studio, RRID:SCR_010970). FASTQ files were processed and mapped with the reference mouse genome (Genome Reference Consortium Build 37) by the Omicsoft Aligner (Omicsoft Sequence Aligner, RRID:SCR_005270). Aligned reads were then quantified into FPKM values at the gene level using the gene model from “RefGene20121217”. Differential gene expression analysis between different groups were performed by generalized linear model using log2 (FPKM+1) values statistical threshold (p < 0.05, q <0.05 and absolute fold change > 1.2).

### Whole genome RNA sequencing of rat OPCs

RNA sequencing of rat OPCs was performed by Genewiz, LLC. (South Plainfield, NJ, USA). Briefly, total RNA was extracted from cell pellet samples using Qiagen RNeasy Plus Universal mini kit following manufacturer’s instructions (Qiagen, Hilden, Germany). Extracted RNA samples were quantified using Qubit 2.0 Fluorometer (Life Technologies, Carlsbad, CA, USA) and RNA integrity was checked using Agilent TapeStation 4200 (Agilent Technologies, Palo Alto, CA, USA).

RNA sequencing libraries were prepared using the NEBNext Ultra RNA Library Prep Kit for Illumina following manufacturer’s instructions (NEB, Ipswich, MA, USA). Briefly, mRNAs were first enriched with Oligo(dT) beads. Enriched mRNAs were fragmented for 15 minutes at 94 °C. First strand and second strand cDNAs were subsequently synthesized. cDNA fragments were end repaired and adenylated at 3’ ends, and universal adapters were ligated to cDNA fragments, followed by index addition and library enrichment by limited-cycle PCR. The sequencing libraries were validated on the Agilent TapeStation (Agilent Technologies, Palo Alto, USA), and quantified by using Qubit 2.0 Fluorometer (Invitrogen, Carlsbad, CA) as well as by quantitative PCR (KAPA Biosystems, Wilmington, USA).

The sequencing libraries were clustered on three lanes of a flowcell. After clustering, the flowcell was loaded on the Illumina HiSeq instrument (4000 or equivalent) according to manufacturer’s instructions. The samples were sequenced using a 2x150bp Paired End (PE) configuration. Image analysis and base calling were conducted by the HiSeq Control Software (HCS). Raw sequence data (.bcl files) generated from Illumina HiSeq was converted into fastq files and de-multiplexed using Illumina’s bcl2fastq 2.17 software (bcl2fastq, RRID:SCR_015058). One mismatch was allowed for index sequence identification.

### RNA extraction, cDNA preparation, and qPCR

RNA isolation from the brain was performed with an RNeasy Lipid Tissue Mini Kit (Qiagen), following manufacturer’s instructions. Briefly, approximately 50 milligrams of brain tissue were homogenized in 1 mL of QIAzol Lysis Reagent with a handheld homogenizer. After separation of the soluble phase by the addition of chloroform and precipitation of nucleotides with 1 volume of 70% ethanol, the samples were pipetted into a RNeasy Mini spin column placed in a 2 mL processing tube and centrifuged for 15 seconds at 8000g. Upon elution with 50 µL RNase free water the RNA was measured and used for cDNA was preparation with the Quantitect Reverse Transcription Kit (Qiagen) according to the instructions provided by the manufacturer. The cDNA was stored at -80℃ until use. RNA isolation from cells was performed using the RNeasy kit (Qiagen) following the manufacturer’s instructions. Samples were eluted in 50μL of RNase free water and RNA was converted to cDNA as above. For qPCR, a master mix solution was prepared with 1X Taqman Fast Advanced Master Mix, a GPR17 probe and an RPL37A probe as the housekeeping control (all from ThermoFisher Scientific). The master mix was aliquoted into a 384 well PCR plate and sample was added to each well. Samples were run in triplicate or as indicated on the QuantStudio 7. (Applied Biosystems)

### GPR17 activity assays

All materials were purchased from ThermoFisher Scientific unless otherwise specified. A CHO Flp-In-T-Rex cell line stably expressing the short isoform of human GPR17 (Blasius, Weber, Lichter, & Ogilvie, 1998; Raport et al., 1996) was generated in house. Cells were maintained at 37⁰C in a humidified incubator enclosing an atmosphere of 95% air/5% CO_2_ in Ham’s F12 media supplemented with GlutaMAX, 10% fetal bovine serum, 100 units/mL of penicillin, 100 µg/mL of streptomycin and 5 µg/mL of Geneticin/Hygromycin. To induce the expression of GPR17, cells were grown to 70% confluency in tissue culture flasks and media was replaced with that supplemented with 10 ng/mL Doxycycline Hyclate (Millipore Sigma, St. Louis, MO, USA) for 24 hours. The media was then removed, and cells rinsed twice with PBS (without Ca^+2^ or Mg^+2^) followed by treatment with 0.05% Trypsin-EDTA to detach cells. The cells were suspended in complete media and centrifuged at 150g for 5 minutes to pellet cells. Supernatant was discarded and the cells rinsed twice with assay buffer for the specific assay to be performed. Finally, the cells were filtered through a 40 µm cell strainer (Corning) and suspended at a density of 7.5 X 10^5^ cells/mL in assay buffer. For the cAMP assay, assay buffer consisted of HBSS supplemented with 20 mM HEPES, 500 μM IBMX (Millipore Sigma) and 0.1% bovine serum albumin, pH 7.4. For agonist/inverse agonist mode, the appropriate amounts of test compound stock solutions prepared in DMSO were pre-stamped into the corresponding well of a white shallow bottom ProxiPlatePlus-384 plate (Perkin Elmer, Waltham, USA) using the ECHO 555 (Labcyte, San Jose, USA). 10 μL of the cell suspension in assay buffer was plated per well using the Multidrop. The plate was briefly centrifuged at 150g, sealed and incubated at room temperature for 1 hour. Following this, 10nL of a 3 mM stock solution of forskolin (Millipore Sigma) was transferred per well using the ECHO 555. The plate was again briefly centrifuged, sealed and incubated for a further hour at room temperature. At the end of the hour, cells were lysed by adding, using the Multidrop (ThermoFisher), 10 μL/well of lysis buffer supplemented with a 1:40 dilution each of the labeled cAMP and correspondingly labeled cAMP antibody pair provided in the cAMP Gi kit (CisBio, Bedford, USA). The plate was briefly centrifuged, sealed and incubated for an hour at room temperature. The HTRF values per well at 665 and 620 nm were obtained by reading the plate in a Pherastar FSX (BMG LabTech, Cary, NC, USA). A ratio of the readings per well at 665/620 nm, multiplied by a factor of 10,000 was used for further calculations. For antagonist mode, the same procedure described above was performed, except for an additional step at the start consisting of preincubation of the test compounds with the cells in the microplate for 30 minutes followed by the addition of an EC80 concentration of the GPR17 agonist MDL29951 (Hennen et al., 2013) or Sanofi’s GPR17 agonist SAR959.

### Cuprizone-induced de/remyelination in C57BL/6 mice. Cuprizone mouse model of demyelination and remyelination

The cuprizone lesion model was performed as previously described (Jurevics et al., 2002; Miron, Kuhlmann, & Antel, 2011) to induce demyelination and follow remyelination events primarily in the corpus callosum region. Briefly, 8- to 10-week-old female and male C57BL/6 mice (7-10 per group, see Diagram 1) were feed a diet containing 0.2% cuprizone (bis-cyclohexanone oxaldihydrazone; Sigma-Aldrich Inc., St. Louis, MO) for up to 5 weeks. Thereafter, mice were fed a normal powder diet for the indicated times to allow spontaneous remyelination to occur. The cuprizone or normal diet were changed twice per week. Food and water were available *ad libitum* and mice were weighed weekly. Cuprizone and control diet animals were sacrificed at the indicated time points, perfused and the brains processed for immunohistochemistry as described elsewhere (Acs, Selak, Komoly, & Kalman, 2013; Anastasiadou et al., 2015; Praet, Guglielmetti, Berneman, Van der Linden, & Ponsaerts, 2014). No randomization was performed to allocate subject in this study.

**Diagram 1.**
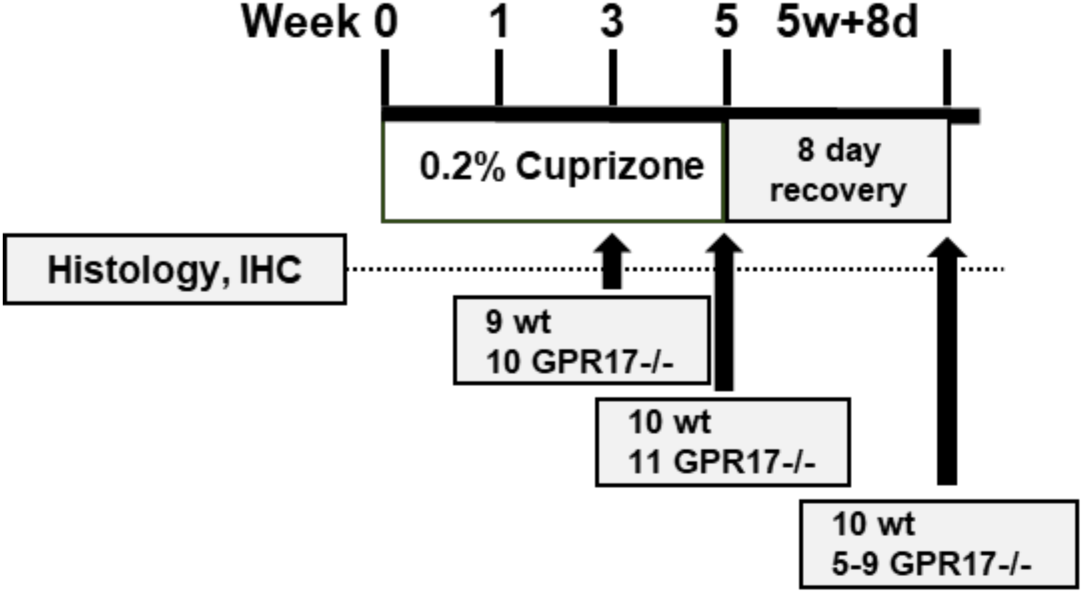
Cuprizone-induced demyelination model used in this study. After 3 and 5 weeks of 0.2% cuprizone administration in the food, mice (wild type and GPR17 -/= and GPR17-/-) were placed in a regular show diet for additional 8 days (black arrows). Histological analysis of myelin protein and mature oligodendrocyte analysis was performed at each of these time points.

### Immunohistochemistry and *in situ* hybridization

Fresh frozen human brain tissue was sectioned at 10 µm with a Microm HM560 cryostat and serial immunohistochemistry was performed with the Roche Ventana Medical Systems DISCOVERY ULTRA platform (Roche, USA) to label tissue sections for the indicated markers. Briefly, slides were wet loaded into the Ventana and exposed to Ventana CC1 for heat induced epitope retrieval. After heat treatment, tissues were blocked using the ChromoMap DAB kit (Roche). Primary antibodies anti-PLP (Abcam, Cat# ab183493, RRID:AB_2833094) and anti-GPR17 (Sigma-Aldrich Cat# HPA029766, RRID:AB_10612114) were diluted in antibody diluent (Roche) and applied to the slides (100 µL/slide). Slides were incubated at 37°C for 1 hour and amplification was then completed with Omni anti-Rabbit HRP (Roche) for 20 minutes. The signal detection was carried out using DAB (Roche) and counterstaining with hematoxylin II and bluing solution (Roche). Finally, slides were removed from the Ventana, dehydrated, and coverslipped with Cytoseal XYL. Slides were dried and then imaged with an Aperio Scanscope AT.

*In situ* hybridization was also performed with the Roche Ventana Medical Systems DISCOVERY ULTRA platform to assess mRNA expression of GPR17 in mouse brain tissue. Briefly, slides were thawed for 2 minutes and placed in 10% neutral buffered formalin for 24 hours then rinsed twice in PBS for 5 minutes and wet loaded into the Ventana. Sections were further permeabilized with cell conditioning for 8 minutes followed by target retrieval at 97°C and protease treatment (16 min at 37°C) (mRNA dewax, mRNA target retrieval VS universal, mRNA protease, all from ACDBIO, USA). Probes were then hybridized for 2 h at 43°C followed by RNAscope amplification. The following RNAscope probes were used in this study: RNAscope 2.5 VS Probe-*Mm-GPR17* (ACDBIO, cat#503199). Final signal detection was carried out using mRNA HRP (RNAscope VS Universal HRP Detection Reagents, ACDBIO) and bluing solution (Roche). Finally, slides were removed from the Ventana, dehydrated, and coverslipped with Cytoseal XYL (Thermo Fisher). Slides were dried and then imaged with an Aperio Scanscope AT.

### Spinal cord lysolecithin de/remyelination

Demyelinated lesions were induced in the dorsal column of the spinal cord of mice as previously described (Najm et al., 2015). Briefly, 9- to 11-week-old C57BL/6 mice were anaesthetized using isoflurane and a T11-T12 laminectomy was performed. 2 μL of a 1% Lysophosphatidylcholine (LPC, Sigma-Aldrich, St. Louis, MO) was infused into the dorsal column at a rate of 0.25μl/min. The animals were euthanized at day 7 after laminectomy (n=7 per group) and the lesion area of the spinal cord processed for histopathological analysis and immunohistochemistry. 20X images acquired by Scanscope AT Turbo (Leica Biosystems Inc., Buffalo Grove, IL) were imported into HALO (Indica Labs, Albuquerque, NM) image analysis software (HALO, RRID:SCR_018350). A region of interest was manually drawn around the dorsal funiculus of the spinal cord and analyzed by tissue classification using the random forest classifier to separate the image into two classes, Lesion and Normal Dorsal tissue. The outputs were area (mm^2^) for region of interest, lesion, and normal dorsal and calculated by using Aperio Scanscope AT and software analysis.

### Statistics

For the comparisons of different treatment groups, column statistical analysis was performed with GraphPad Prism software (GraphPad Prism, RRID:SCR_002798). Statistical analysis of experiments performed with mice (n=9-11 animals/group) was carried out using one-way ANOVA for grouped comparisons and Student’s *t*-test for comparisons between two groups. p values are indicated by * ≤ 0.05, ** ≤ 0.01, *** ≤ 0.001, and **** ≤ 0.0001. For *in vivo* experimentation, the experimenter was unaware of the animal groupings during imaging and statistical analyses.

## Results

### GPR17 expression levels peak at early stages of developmental OPC maturation and are elevated in demyelinated tissue

OPCs (A2B5^+^/platelet-derived growth factor alpha^+^, PDGFRα) exit the cell cycle to differentiate into multi-branched cells, extending long processes, forming complex membranous structures and targeting axons for subsequent membrane wrapping and myelin compaction (Bauman, & Pham-Dinh, 2001). These dramatic morphological transformations are driven by a changing genetic signature geared to the production of structural myelin proteins such as MBP (myelin basic protein), PLP (proteolipid protein) and MOG (myelin oligodendrocyte glycoprotein) among others. At the same time, early OPC genes like PDGFRα and NG2 (chondroitin sulfate proteoglycan 4) are downregulated. We have detected GPR17 mRNA expression significantly elevated in early OPC stages, peaking after 24-48h *in vitro*, then decreasing to low levels as the cells differentiate towards pre-myelinating oligodendrocytes (Figure 1, A). Correspondingly, early protein expression was detected by immunocytochemistry in branched OPCs, positive for the progenitor marker NKX2.2 (B). Maturing oligodendrocyte lineage cells expressing MBP showed no GPR17 immunoreactivity (Figure 1, B, C). This temporal pattern of GPR17 expression in relation to myelin protein dynamics supports the hypothesis of GPR17 influencing developmental maturation processes and confirms previous reports by other laboratories on the pattern of GPR17 expression in OPCs (Ceruti et al., 2011; Chen et al., 2009; Ciana et al., 2006; Lecca et al., 2008).

**Figure 1.**
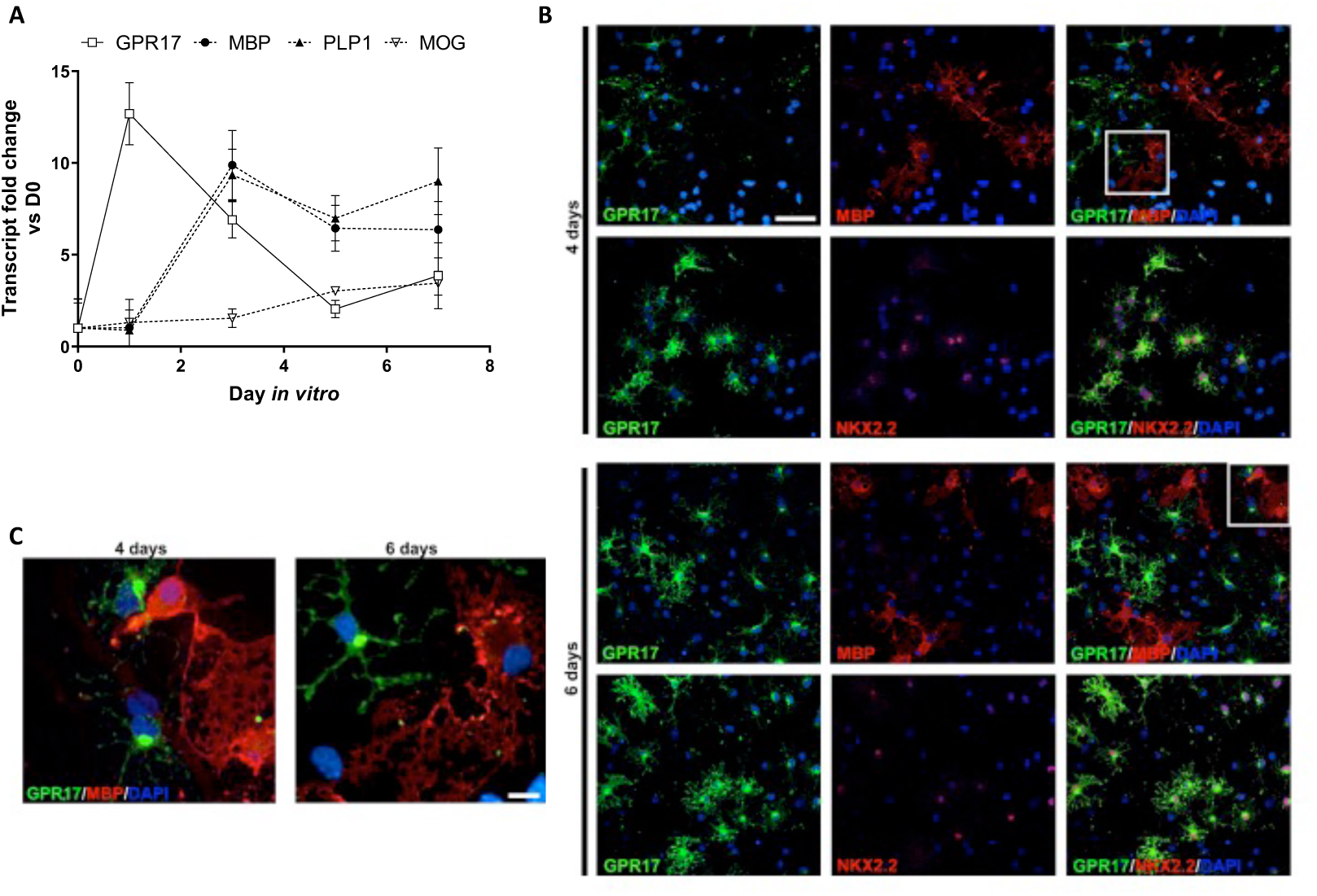
GPR17 expression is high in early oligodendrocyte precursors *in vitro*. Cultured rat OPCs showed high levels of GPR17 transcript at early stages of maturation, returning to baseline levels as the cells transitioned into pre-myelinating oligodendrocytes (A, open squares). Increased transcript levels for the myelin proteins, myelin basic protein (MBP; A, black circles), proteolipid protein (PLP; A, black triangles) and myelin oligodendrocyte glycoprotein (MOG; A, open triangles) demonstrates OPC maturation. GPR17 transcript expression correlated with protein levels detectable by immunocytochemistry in NKX2.2^+^ cells, a marker of immature progenitors, showing morphological features of early OPCs (B, lower panels in 4- and 6-day images; C). Upon maturation into MBP^+^ cells and appearance of membranous structures, the expression of GPR17 was downregulated to undetectable levels (B, top panels in 4- and 6-day images; C). RNAseq data is the result of RNA isolation from triplicate wells per time point (see material and methods for data analysis details).

The presence of OPCs in and around MS lesions has been extensively documented (Boyd, Zhang, & Williams, 2013; Kuhlmann et al., 2008). A number of these progenitors (NG2^+^/PDGFRα^+^) appear to extend processes and branches but fail to fully mature into myelinating oligodendrocytes. We analyzed GPR17 expression, using immunohistochemistry, in postmortem tissue from a relapsing remitting MS patient. Cells positive for GPR17 had a morphological appearance reminiscent of OPCs and, while a few GPR17^+^ cells were found within the lesion (PLP^-^area in Figure 2, A), a significant congregation of GPR17^+^ was observed primarily in peripheral areas where demyelination was less severe or absent (PLP^+^ area in Figure 2, A). This pattern of GPR17 expression in MS tissue was also observed in several samples from secondary and primary progressive patients (data not shown). We analyzed GPR17 mRNA expression levels in MS samples (relapsing remitting, secondary progressive and primary progressive) and found GPR17 expression to be slightly higher in MS samples than in non-lesion areas. However, there was high variability among samples and these differences were not statistically significant (Figure 2, B).

**Figure 2.**
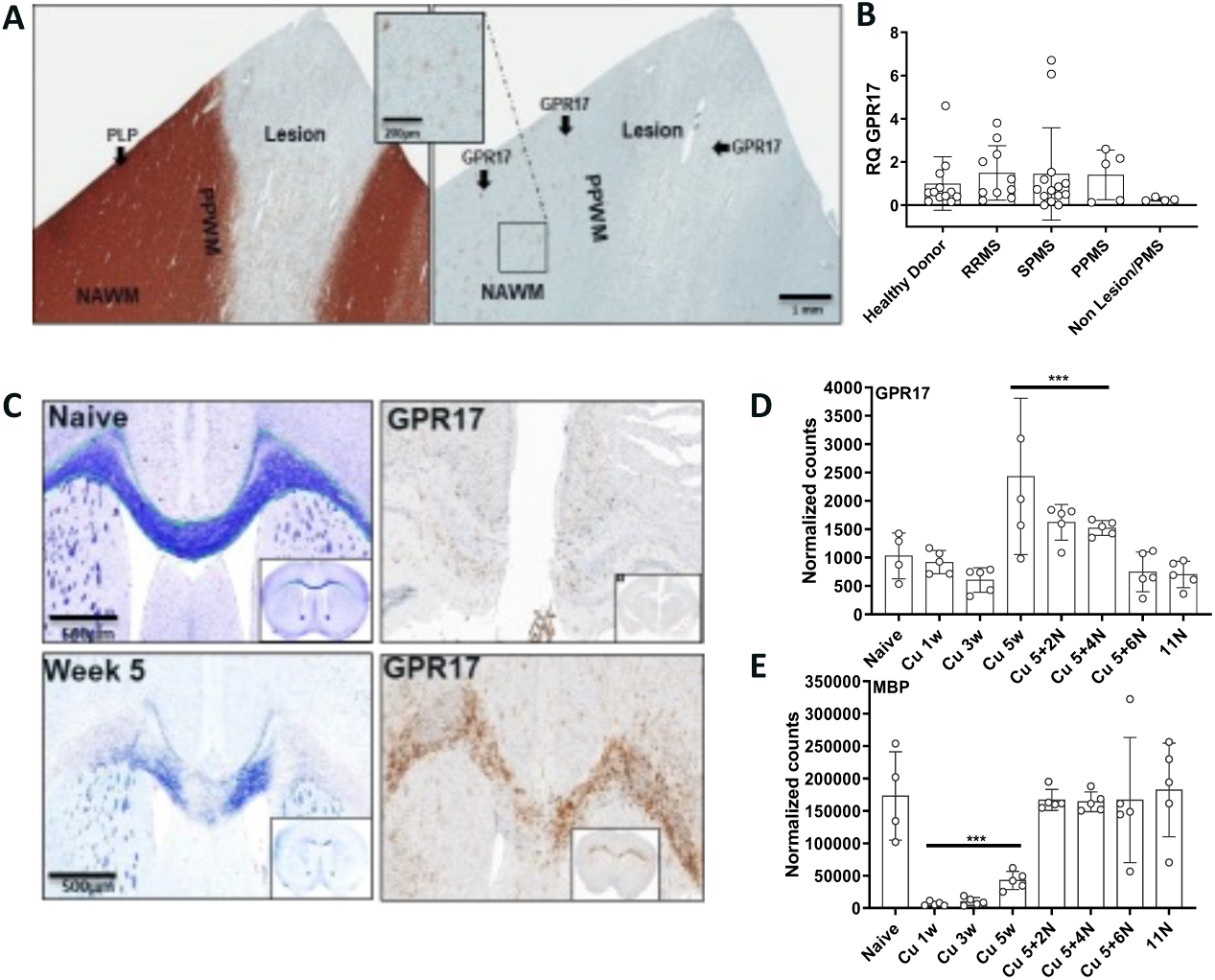
GPR17 expression is increased in demyelinated lesions. Multiple sclerosis demyelinated lesions were identified in postmortem brain tissue by lack of proteolipid protein immunostaining (PLP, A, left panel). GPR17 positive cells showing branching morphology, were detected in peri-lesion (PPWM) and normal appearing white matter areas (NAWM; A, right panel). Fewer GPR17 positive cells were detected in extensively demyelinated tissue. GPR17 transcript levels were measured by real-time PCR in tissue containing a lesion from postmortem patients with relapsing (RRMS) or progressive pathologies (B, SPMS, PPMS). No significant differences in GPR17 levels were detected among the samples used in this test (RQ: relative expression to RPL37a transcript levels used as a housekeeping, reference control). C-D shows C57/Bl6 mouse brain tissue where chemical demyelination was induced in response to up to 5 weeks of cuprizone diet, followed by up to 6 weeks of normal diet to allow for spontaneous remyelination to occur. Demyelination was measured by global lipidic staining with Luxol Fast Blue in the corpus callosum at week 5 of cuprizone diet (C, left panels). GPR17 immunostaining showed abundant OPCs in demyelinated areas at 5 weeks of cuprizone (C, right panels). Coronal mouse brain sections containing the ventral region of the corpus callosum were processed for laser microdissection and RNA isolation at the time points indicated in D and E. RNA sequencing and analysis of transcripts of interest is expressed as normalized counts from 4- 5 animals in each condition. Myelin basic protein (MBP) transcript levels were significantly reduced soon after the beginning of cuprizone ingestion (E, cuprizone 1 through 5 weeks, Cu 1w-5w), returning to naïve levels after two weeks of normal diet (E, Cu 5 + 2N) until the end of the experiment at week 11 (Cu 5 + 6N). GPR17 RNA levels were significantly elevated at the peak of remyelination (Cu 5 + 2N – Cu 5 + 4N). *** p<0.01 according to one-way ANOVA.

We next aimed to characterize the expression pattern of the receptor in a demyelination/remyelination animal model. To this end, we used demyelinated tissue in mice fed the neurotoxicant cuprizone, which has been extensively characterized previously (Hiremath et al., 1998; Matsushima & Morell, 2001; Praet et al., 2014), and involves oligodendrocyte (OL) cell death and a continuous maturation of endogenous OPCs, actively remyelinating damaged tissue. GPR17 expression was determined by RNA sequencing of micro-dissected corpus callosum areas during cuprizone-induced demyelination (0.2% in chow, weeks 1-5) and remyelination following toxicant withdrawal (normal diet, weeks 5-11). Receptor mRNA expression was significantly high upon returning the animals to normal diet, marking the beginning of remyelination and remaining elevated until myelin content determined by MBP expression, was back to normal levels (Figure 2, D, E). Correspondingly, elevated GPR17 protein immunoreactivity was detected in the corpus callosum of 5-week cuprizone-fed mice at the peak of demyelination (Figure 2, C). These data show elevated GPR17 expression in cells undergoing maturation which recapitulates the pattern observed during developmental OL formation.

### GPR17 deletion results in a gene signature with minor impact on myelin gene expression

It has been previously described that GPR17 knockout mice (GPR17^-/-^) show an early onset of developmental myelination and the opposite, i.e. hypomyelination, when the receptor is overexpressed in OPCs under the CNPase (2’,3’-cyclic nucleotide-3’-phosphodiesterase) promoter (Chen et al., 2009). With the goal of further understanding the impact of GPR17 expression on myelin formation *in vivo*, we implemented a GPR17^-/-^ mouse previously characterized (Mastaitis et al., 2015) and utilized it for histopathological analysis of myelin dynamics as well as to generate OPC cultures for *in vitro* maturation assays. Receptor deletion was confirmed in GPR17^-/-^ mice by both *in situ* hybridization (ISH) and immunohistochemistry (IHC), in postnatal day 3 (p3) pups in comparison to wild type (Wt) litter mates (Figure 3, A). At this time point, we observed no differences in MBP immunoreactivity in tissue sections from Wt vs GPR17^-/-^ (data not shown) and although a few myelinated axons were detected at the ultrastructural level by electron microscopy, the variability was extremely high among samples, making quantification unachievable (Figure 3, A bottom panels).

**Figure 3.**
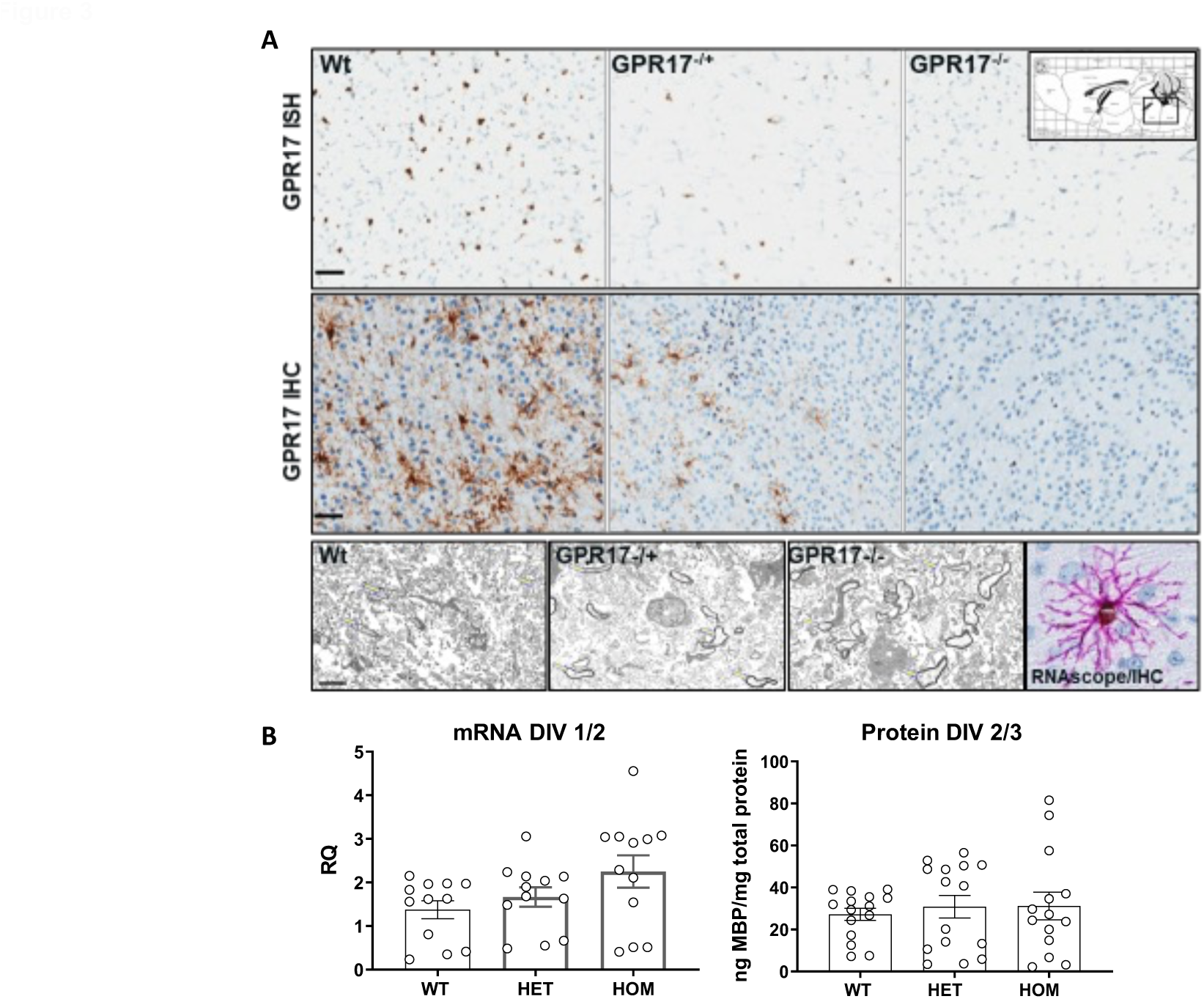
Absence of GPR17 results in a minor enhancement of myelin production during development *in vivo* and *in vitro* OPC maturation. Postnatal day 3 mouse brain tissue was used to characterize the genetic GPR17 knockout (Hom) at the mRNA level by *in situ* hybridization (ISH) and at the protein levels by immunohistochemistry (IHC), in comparison to heterozygous (Het) and wild type (Wt) littermates. GPR17 mRNA and protein were widely expressed throughout the brain in Wt mice, reduced in Het and completely absent in Hom brains (A, top and middle panels). GPR17 was observed in branched cells with OPC morphology, localized in cell processes and neurites as determined by IHC (A, middle panel) and RNAscope (A, bottom, right panel). A few and sparse, thinly myelinated axons were detected in all conditions by means of electron microscopy (A, bottom panels). Studies of cultured OPCs isolated from these mice showed a minor, non-statistically significant increase (unpaired Student’s t-test) in MBP mRNA expression as well as MBP protein expression in cells obtained from GPR17 knockout (Hom) as compared to Het and Wt littermates (B). RQ: relative expression; IHC: immunocytochemistry; ISH: *in situ* hybridization; Wt: wild type; Het: Heterozygous; Hom: Homozygous; DIV: day *in vitro*. ISH and IHC bars: 50μm; EM bar: 2μm.

OPC cultures were then generated to facilitate the study of receptor deletion in *in vitro* maturation assays as well as gene expression and signal pathway analysis. OPCs from GPR17^-/-^ (homozygous and heterozygous) and Wt pups (p1-3) were maintained *in vitro* in low growth factor media, in order to stimulate their spontaneous maturation. Then, after RNA and protein extraction, MBP expression was measured as an indication of the level of differentiation. We found no statistically significant differences between GPR17^-/-^ versus Wt OPCs for MBP expression at both the RNA and protein levels (Figure 3, B). These data indicate that GPR17 has a minor, transitory effect on OPC maturation *in vitro* and suggest that only a small population of cells are susceptible to changes driven by receptor expression and function, resulting in a minor enhancement of global developmental myelination *in vivo*.

In order to investigate in depth the correlations between GPR17 and OPC/OL gene expression, we performed whole genome RNA sequencing in cultured mouse OPCs, comparing GPR17^-/-^ versus Wt. Purified mouse OPCs from GPR17^-/-^ and Wt pups at postnatal day 1-3 were cultured for different time points as indicated, RNA isolated and gene expression determined and analyzed by NGS methodologies. As observed in Figure 4, GPR17 deletion was confirmed in GPR17^-/-^ versus high levels detected in the Wt cells where its expression increases from D0 to D1 showing a sharp decrease by D3 (Figure 4 B, top panel). However, these clear changes in GPR17 expression, also observed in rat OPCs (Figure 1, A), did not correspond to elevated levels of myelin gene expression. We observed a minor increase in MBP mRNA levels in GPR17^-/-^ cells only at D0 while PLP mRNA levels trail behind those in Wt cells at D1 and D3, showing no difference at D0 (Figure 4, B).

**Figure 4.**
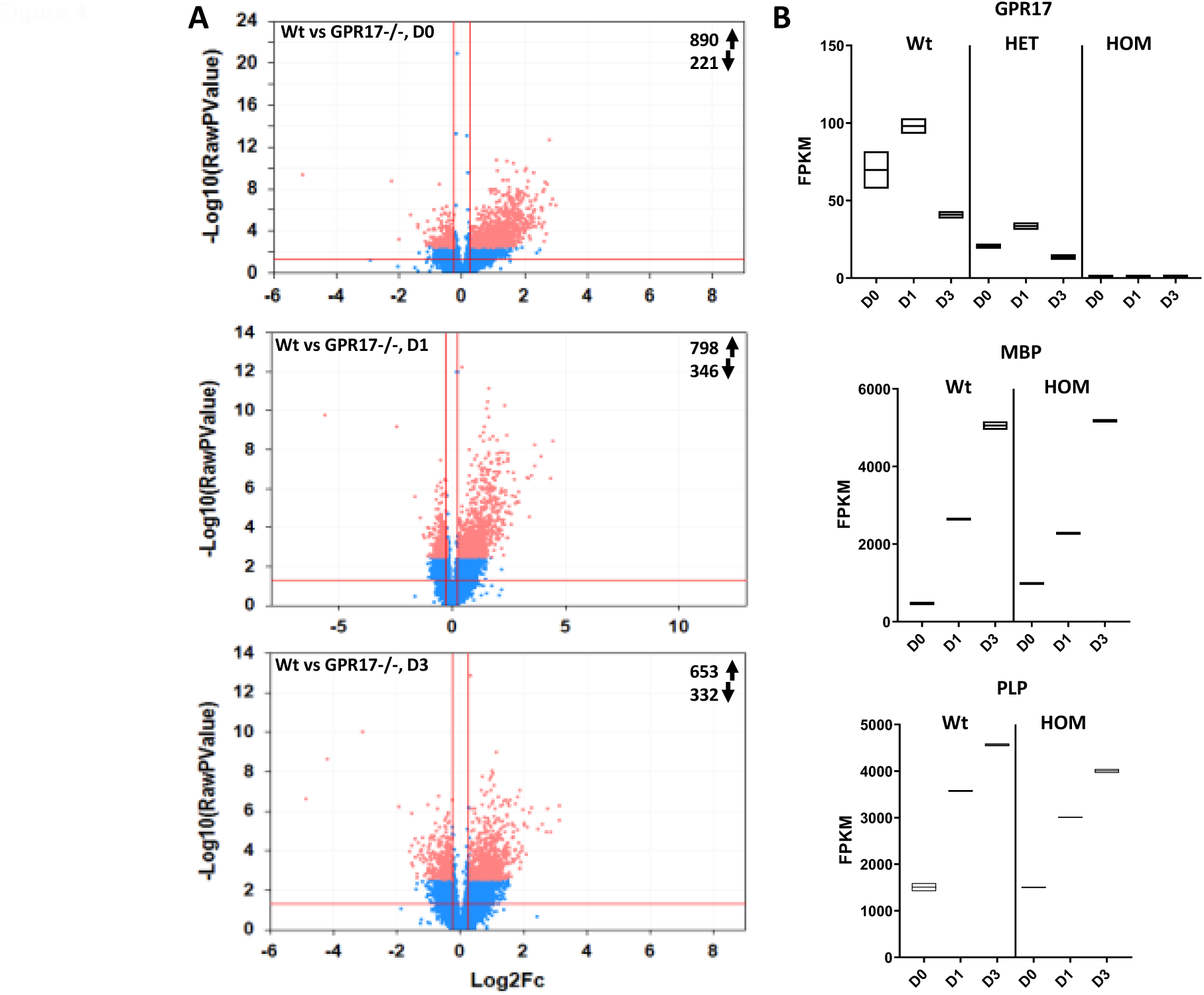
GPR17 knockout OPCs present minor changes in myelin gene transcripts. Total RNA was isolated from cultured OPCs generated from wild type (Wt) or GPR17 knockout (GPR17^-/-^, Het and Hom) newborn pups at the day of plating (D0), and after 1 (D1) and 3 (D3) days *in vitro*. Whole genome mRNA sequencing was performed, and transcript expression analysis carried out. At all timepoints studied, differentially expressed genes (DEGs) were identified between Wt and GPR17^-/-^ (A, arrows indicate number of genes being upregulated and downregulated at each time point). GPR17 knockout was confirmed by the complete absence of transcript in the Hom cells (B, top panel). In Wt OPCs, GPR17 is highly expressed at early time points, decreasing significantly at D3. There were no major differences in myelin gene expression (MBP and PLP are shown) between Wt and GPR17^-/-^ (Hom, B middle and bottom panels). Data are presented as raw *p* values (Log10) in the volcano plots and as fragments for kilobase of transcript per million (FPKM) in the histograms.

Although only minor changes were observed in myelin gene expression upon GPR17 deletion, we were able to identify a gene signature associated with GPR17 absence in mouse OPCs. At D0, the expression of 221 genes was downregulated and 890 upregulated in cells lacking GPR17 (Figure 4, A, top panel). As the cells matured, the number of downregulated genes was 332 versus 653 trending up by D3 (Figure 4, A, bottom panel). Upon clustering mRNAs by Gene Ontology, it was evident that GPR17 deletion correlates with changes in expression of genes peripherally involved in myelin dynamics, e.g. axonal guidance and gap junction signaling. Also in the list of upregulated genes at D1 showing a greater than 2-fold change in GPR17^-/-^ cells versus Wt, appeared several OPC and myelin synthesis and regulation related proteins, including myelin glycoprotein (Mgp, +5.9), matrix metalloproteinase-2 and 3 (Mmp2, Mmp3, *+*5.4 and *+*20.2 respectively), glypican-4 (GPC4, *+*4.6) indicating an indirect or secondary regulatory influence of GPR17 expression in the myelination process (Tables 1-3).

**Table 1.**
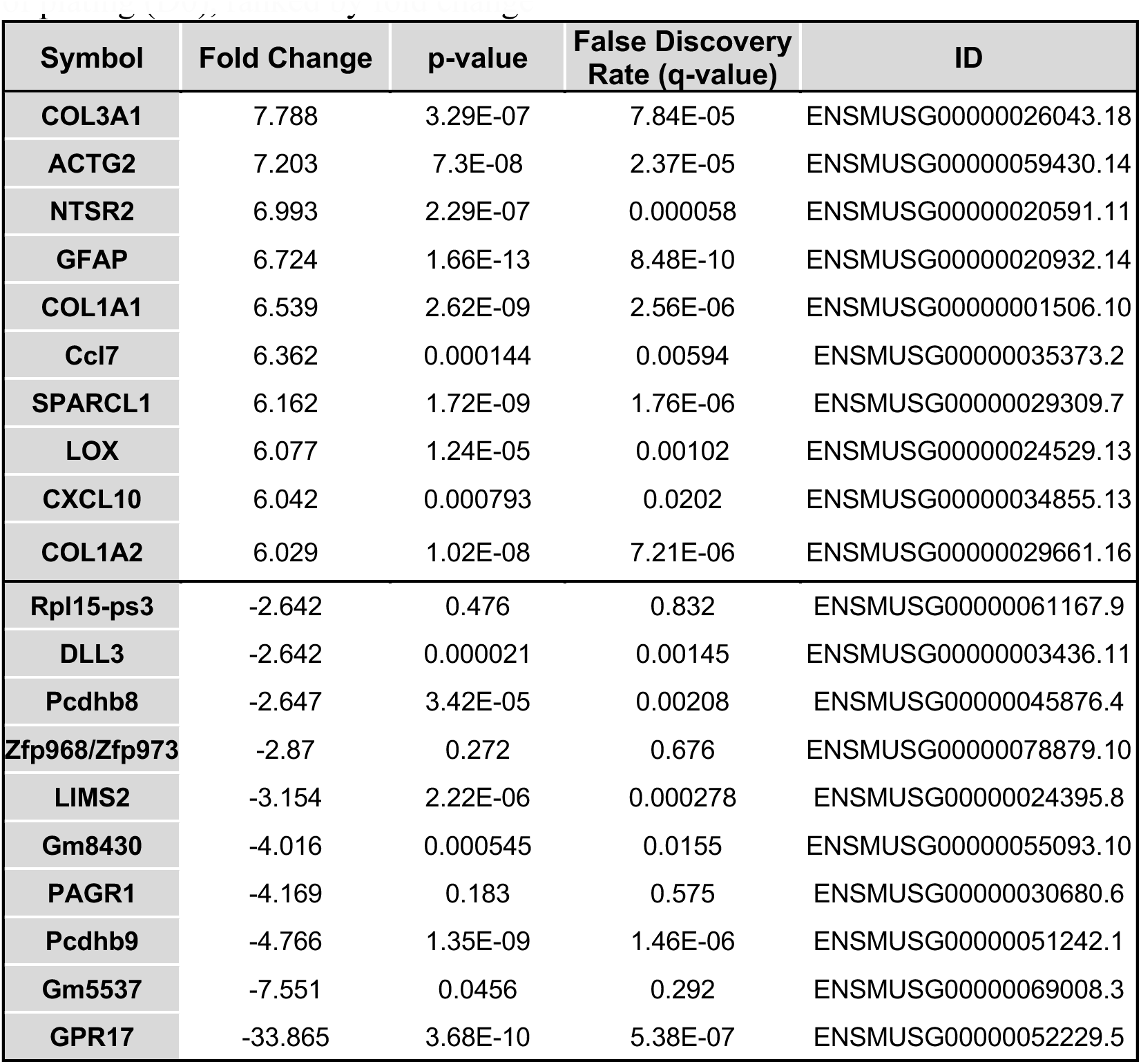
Example of significantly differentially expressed genes in GPR17-/- vs. Wt cells at day of plating (D0), ranked by fold change.

**Table 2.**
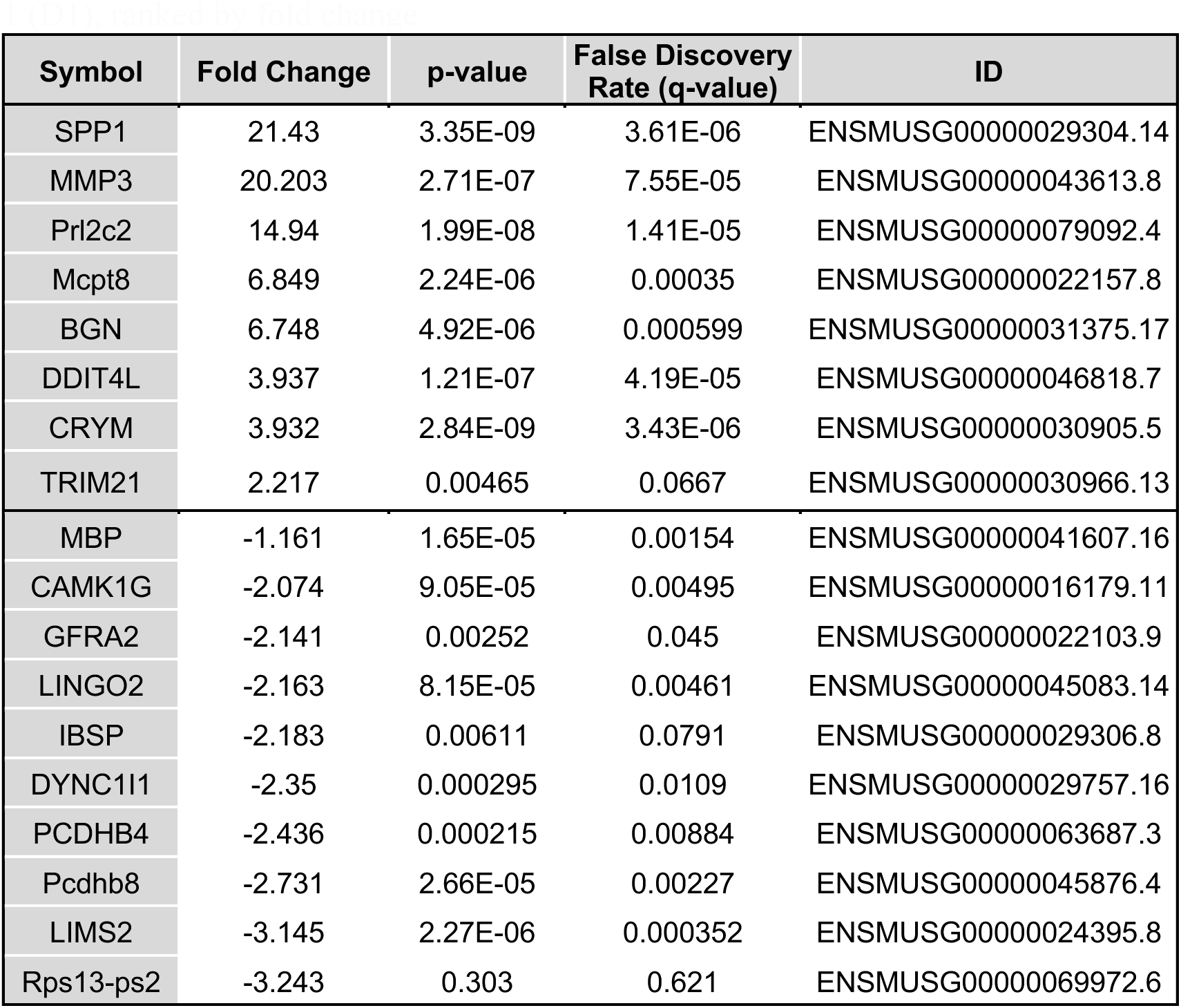
Example of significantly differentially expressed genes in GPR17-/- vs. Wt cells at day 1 (D1), ranked by fold change.

**Table 3.** Example of significantly differentially expressed genes in GPR17-/- vs. Wt cells at day 3 (D3), ranked by fold change.

| Symbol | Fold Change | p-value | False Discovery Rate (q-value) | ID |
| --- | --- | --- | --- | --- |
| Saa3 | 8.598 | 2.57E-06 | 0.000864 | ENSMUSG00000040026.8 |
| MMP13 | 8.568 | 4.76E-07 | 0.000296 | ENSMUSG00000050578.10 |
| HCLS1 | 3.972 | 4.45E-06 | 0.00114 | ENSMUSG00000022831.14 |
| CXCL6 | 3.931 | 3.47E-06 | 0.000954 | ENSMUSG00000029371.7 |
| RET | 2.705 | 0.00012 | 0.00875 | ENSMUSG00000030110.13 |
| DCN | 2.699 | 0.000243 | 0.0123 | ENSMUSG00000019929.16 |
| GYPC | 2.698 | 0.0016 | 0.038 | ENSMUSG00000090523.2 |
| PXMP2 | 2.287 | 0.000954 | 0.0286 | ENSMUSG00000029499.11 |
| STMN2 | 2.287 | 0.00362 | 0.0608 | ENSMUSG00000027500.10 |
| MBP | 1.025 | 0.21 | 0.551 | ENSMUSG00000041607.16 |
| MT-RNR1 | -2.049 | 0.000903 | 0.0277 | ENSMUSG00000064337.1 |
| SPOCK3 | -2.051 | 0.0133 | 0.132 | ENSMUSG00000054162.15 |
| PLPPR1 | -2.066 | 4.15E-07 | 0.00029 | ENSMUSG00000063446.4 |
| TCERG1L | -2.304 | 1.98E-05 | 0.00295 | ENSMUSG00000091002.2 |
| EFNB2 | -2.329 | 0.000143 | 0.00957 | ENSMUSG00000001300.16 |
| SLC35G2 | -2.339 | 0.00321 | 0.0572 | ENSMUSG00000070287.3 |
| Rps27/rt | -2.374 | 0.248 | 0.594 | ENSMUSG00000050621.7 |
| Gm10401 | -3.675 | 0.0731 | 0.342 | ENSMUSG00000072693.2 |
| LIMS2 | -3.886 | 5.39E-07 | 0.000318 | ENSMUSG00000024395.8 |
| Pcdhb9 | -8.497 | 8.17E-11 | 8.37E-07 | ENSMUSG00000051242.1 |

### Highly selective and potent GPR17 activity modulators have a moderate effect on OPC maturation

In order to identify small molecules able to modulate receptor activity, we developed and optimized GPR17 cellular assays in both CHO cells stably expressing the short form of the human GPR17 in a doxycycline-inducible manner, and primary rat OPC cultures. We analyzed GPR17-mediated intracellular second messenger (cAMP) accumulation and mobilization of Ca^+2^ via Gαi or Gq signaling modes, respectively. Using these assays, an extensive small molecule screen campaign was carried out through Sanofi chemical libraries. This effort was complemented by medicinal chemistry and structure-activity relationship methodologies which allowed us to identify several specific and potent GPR17 agonists. Proprietary compounds were tested along with chemicals previously characterized elsewhere as GPR17 modulators such as the agonist MDL29951 (Hennen et al., 2013), the antagonist HAMI3379 (Merten et al., 2018) and the cysteinyl leukotriene receptor modulators pranlukast and montelukast, previously described as GPR17 active chemicals (Ciana et al., 2006; Hennen et al., 2013; Ou et al., 2016).

One of the compounds identified in our small molecule screen, SAR959, displayed a potent ability to decrease cAMP production induced by the adenylyl cyclase activator forskolin (agonist mode) with EC50 values in the picomolar range (EC50: 6.5^-10^), approximately 169 times more potent than MDL29951 (EC50: 1.1^-7^, Table 4). Using this potent agonistic activity as a baseline, we determined that pranlukast, HAMI3379, and several Sanofi scaffolds showed antagonistic activity in the low micromolar to nanomolar ranges (Table 4), confirming previously described modulatory activity of the receptor by these compounds (Ciana et al., 2006; Merten et al., 2018). Chemicals showing activity in the engineered CHO cells were then tested in OPC cultures where, due to low activity in the Gαi mode, the most reliable method for determining GPR17 activity was measurements of Ca^+2^ mobilization. Treatment of OPCs with SAR959 effectively mobilized intracellular Ca^+2^ (EC50: 5.2^-8^), 29-fold more potently than MDL29951 (Table 5). In addition, a structural analog of SAR959 termed SAR955, was unable to induce changes in either cAMP or Ca^+2^ in either cell type. We used this compound as a negative control in subsequent assays, effectively validating the methods used (Table 5).

**Table 4.** Compounds showing functional activity in CHO cells over-expressing GPR17.

| **Agonists** | **EC50 [M]** | **Antagonists** | **IC50 [M]** |
| --- | --- | --- | --- |
| MDL29951 | 1.14E-07 | BayCystLT | 5.26E-05 |
| SAR455 | 3.70E-08 | Pranlukast | 2.85E-05 |
| SAR491 | 2.50E-08 | Montelukast | 1.27E-05 |
| SAR511 | 1.70E-09 | HAMI3379 | 7.97E-06 |
| SAR467 | 2.50E-09 | SAR592 | 3.70E-06 |
| SAR959 | 6.50E-10 | SAR901 | 6.05E-07 |

**Table 5.** Comparison of GPR17 activity in CHO cells and primary rat OPCs using the cAMP accumulation (Gi mode) and calcium mobilization (Gq) methods.

| **Assay** | **SAR959**<br>**EC50 [M]** | **MDL29951**<br>**EC50 [M]** | **SAR955**<br>**EC50 [M]** |
| --- | --- | --- | --- |
| **CHO cAMP Gi Agonist** | 6.50E-10 | 1.10E-07 | ND |
| **CHO FLIPR Ca Gq Agonist** | 3.00E-09 | 6.00E-07 | ND |
| **Rat OPC cAMP Gi Agonist** | ND | ND | ND |
| **Rat OPC FLIPR Ca Gq Agonist** | 5.21E-08 | 1.50E-06 | ND |

With proven compound modulation of GPR17 activity in a cell line and in primary OPCs, we then measured functional activity of potent compounds in cultured OPCs in terms of MBP production (OPC maturation). Responses to strong GPR17 agonists was variable as SAR959 (EC50: 5.2^-8^) showed no significant effect on OPCs, while SAR467 and SAR455 (EC50: 2.5x10^-9^ and 3.7x10^-8^ respectively) induced a 2-2.5-fold increase in MBP production (Figure 5 B, left panel, Table 4). Notice the lack of concentration-response effects for these OPC active compounds. On the other hand, antagonistic compounds such as montelukast and SAR592 (IC50:1.27x10^-05^ and 3.7x10-^06^ respectively) induced a significant maturation effect at the lowest concentrations used (Figure 5 C, Table 4). Pranlukast and HAMI3379, previously described to enhance OPC maturation (Ciana et al., 2006; Merten et al., 2018), failed to induce such an effect in our cultures at the concentrations tested. Noticeably, regulation of GPR17 activity in OPCs had a minor effect in cell maturation when compared to clemastine or T3, modulators of signaling pathways independent of GPR17 and used as internal controls in the present study. As expected SAR955 used as a negative control of GPR17 activity, showed no effect on OPC maturation (Figure 5, top right graph).

**Figure 5.**
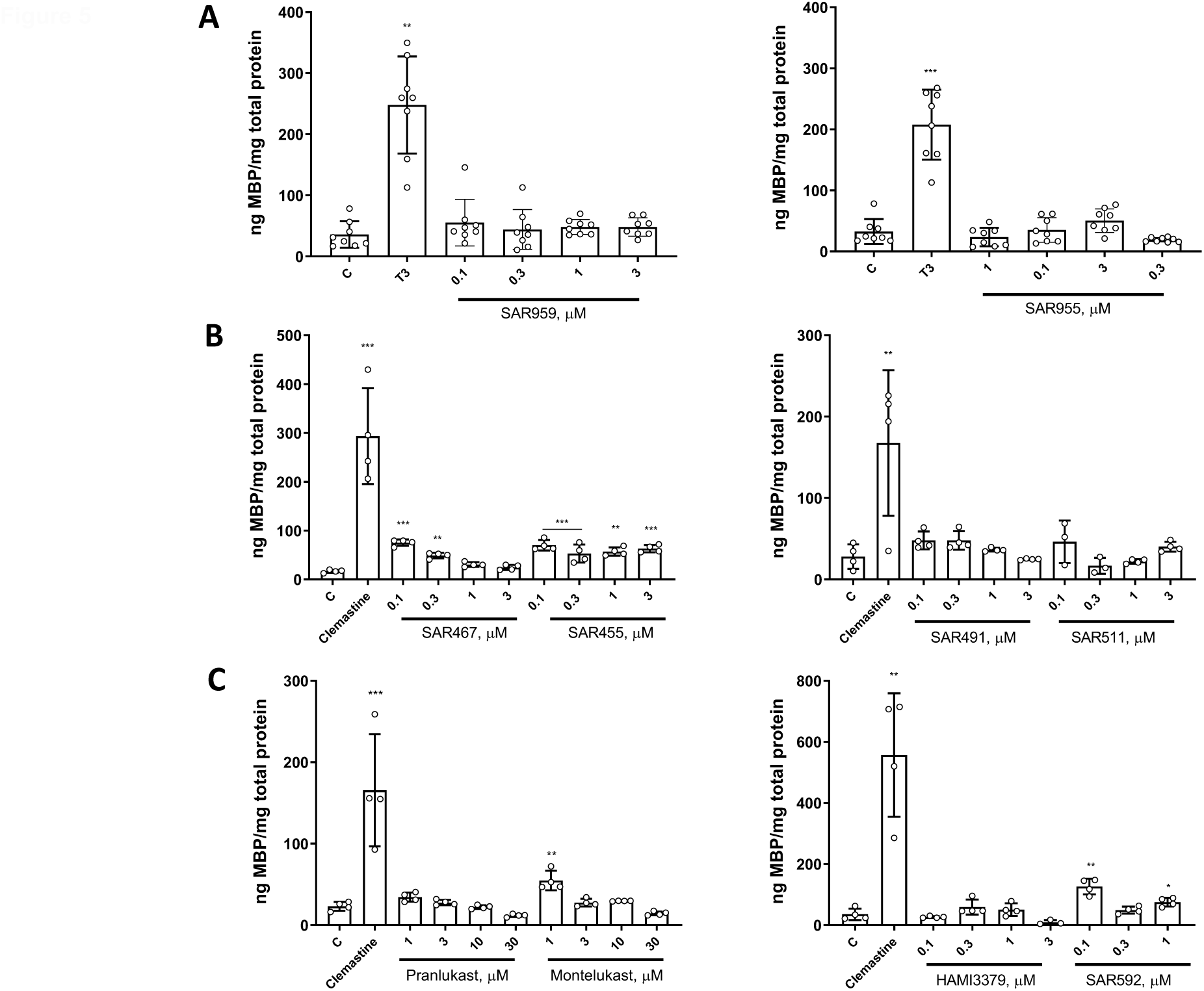
Small molecule GPR17 ligands induce a slight increase in OPC maturation. Compound modulators of GPR17 activity induced a mild enhancement of OPC maturation as determined by the levels of myelin basic protein (MBP). From a series of agonistic compounds (A and B rows), SAR467 and SAR455 induced approximately a 2-fold increase in OPC maturation at some of the concentrations tested, without showing a concentration-response effect (B, left graph). SAR959 (A, left graph), SAR491 and SAR511 (B, right graph) had no effect on MBP production in cultured OPCs. Antagonistic compounds montelukast and SAR592 (C) showed a measurable maturation effect at variable concentrations, while HAMI3379 and pranlukast failed to induce OPC maturation (C). Data are presented as mean ± SD of quadruplicate wells in each treatment. Plotted data are n=2 (top panels) or representative result of 2 independent experiments (B, C). ***: p<0.001; **: p<0.01; *: p<0.05, according to Student’s t test.

### Remyelination is not enhanced in GPR17^-/-^ mice after demyelination induced by cuprizone ingestion or by focal lysolecithin injections

Our *in vitro* experimentation demonstrates that modulation of GPR17 activity by selective and potent small molecules resulted in a minor effect on OPC maturation. To a large extent, OPCs in culture recapitulate the events of developmental differentiation which could differ from endogenous maturation of tissue resident adult OPCs, the cells responsible for maintaining and repairing damaged myelin in demyelinating conditions. To address the possibility of GPR17 activity influencing adult OPC maturation and remyelination, we explored the effect of GPR17 absence on two animal models of de/remyelination. First, we used chemically induced demyelination with focal injections of lysophosphatidylcholine (LPC) which acts as a bio-detergent causing a diffused lesion that spontaneously remyelinates (Hall, 1972; Jeffrey, & Blakemore, 1995). As shown in Figure 6, we observed remyelinating lesions in both wild type (WT) and GPR17 knockout (GPR17KO) after SC dorsal injections of LPC. However, there were no noticeable differences between conditions or enhanced myelin recovery in GPR17KO tissue at 8 days post-lesion. In these tissues, myelin content and the extent of the lesions were determined by both myelin oligodendrocyte glycoprotein (MOG) immunostaining and global lipidic staining with ECR (Eriochrome cyanine R, Figure 6).

**Figure 6.**
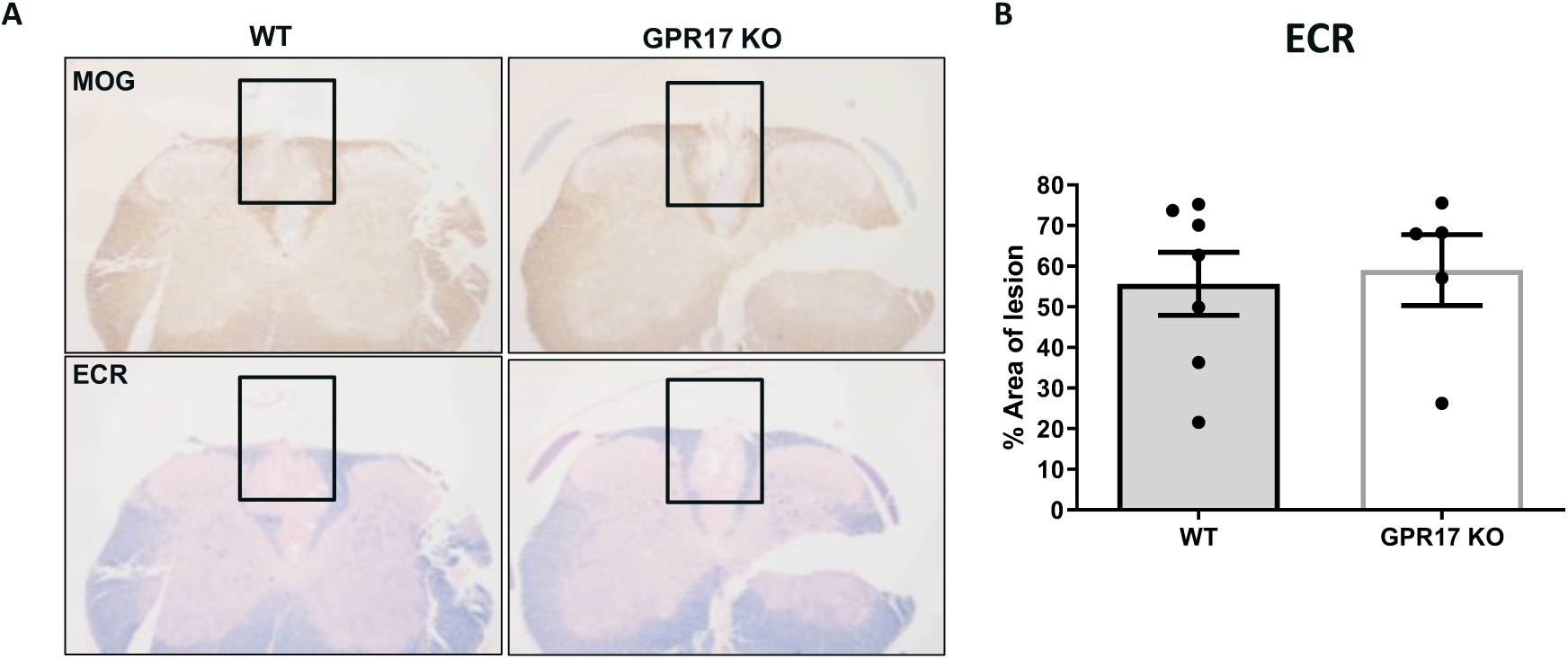
GPR17 absence has no impact on remyelination after focal spinal cord LPC-demyelination insult. Focal demyelination in wild type (WT) or GPR17 knockout (GPR17KO) mouse lumbar spinal cord was induced by injecting 2μL of lysophosphatidylcholine (LPC, lysolecithin) and measured 4- and 8-days post-lesion by Erichrome cyanine R staining (ECR, A, bottom panels) and immunodetection of myelin oligodendrocyte glycoprotein (MOG, A, top panels). Area covered by ECR was determined by software processing of regional densitometry. No significant differences were observed in the rate of remyelination observed between WT and GPR17KO (B). Data plotted in B are the result of analyzing spinal cord sections from 5-7 animals/condition

We then implemented a broader demyelination insult in the brain by means of the neurotoxicant cuprizone, administered in the food (0.2% in solid chow for 5 weeks) as widely described in the literature (Hiremath et al., 1998; Matsushima & Morell, 2001; Praet et al., 2014). In this model, spontaneous remyelination ensues upon returning the mice to normal diet for an additional 4 to 8 days, as a result of a rapid maturation of endogenous, resident OPCs. Once again, we did not observe an enhancement of myelin recovery in GPR17 knockout (HET or HOM) versus wild type (WT) mice as determined by global lipid staining with Luxol Fast Blue (LFB) or MOG and GSTpi immunodetection for myelin and number of mature oligodendrocytes, respectively (Figure 7, A-C).

**Figure 7.**
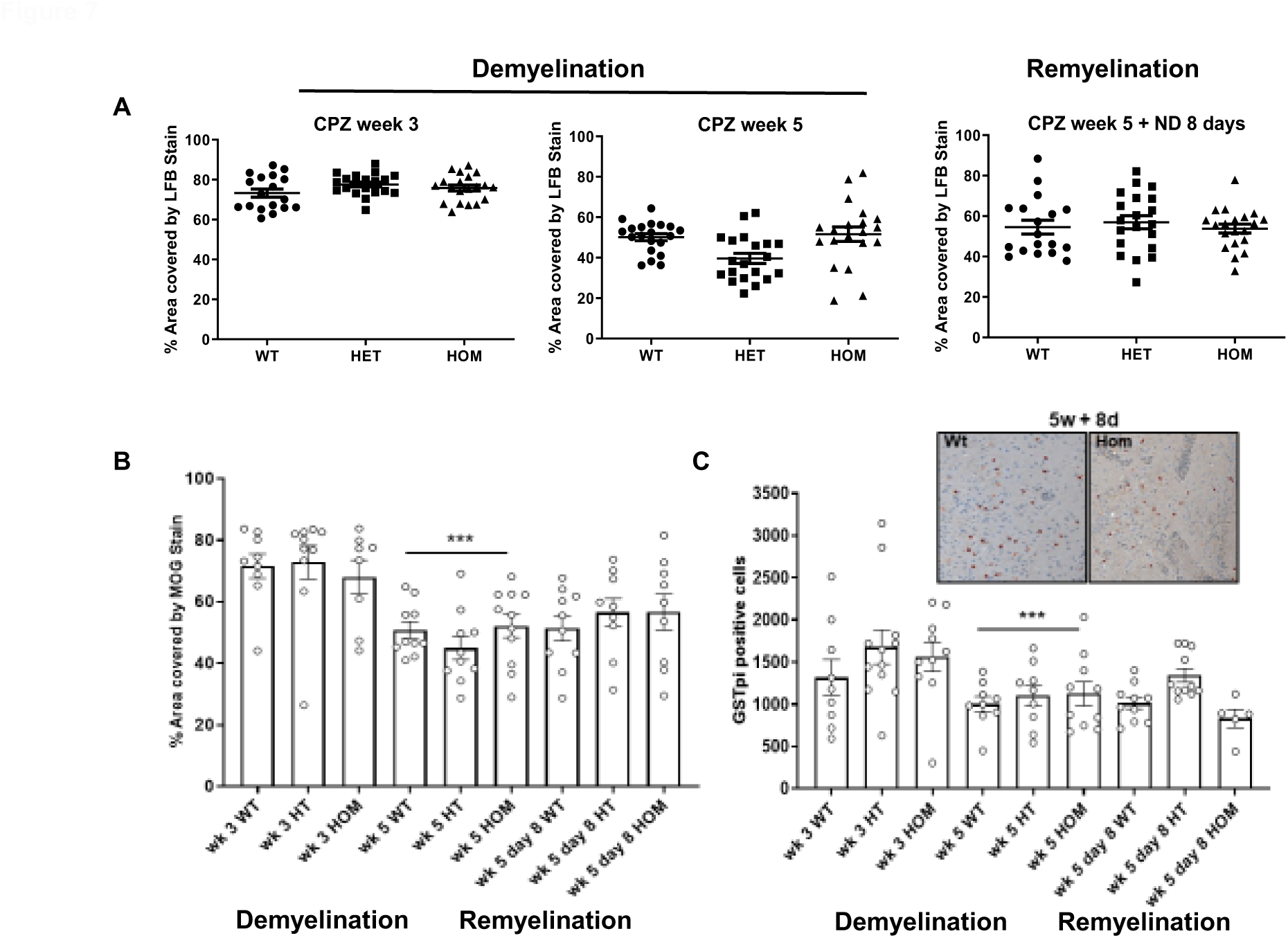
GPR17 knockout does not protect from demyelination nor enhance spontaneous remyelination in the cuprizone mouse model of chemical demyelination. Wild type (WT n=9-10) or GPR17 knockout mice (HET, HOM n=10-11) were fed a diet containing 0.2% of the copper chelator cuprizone (CPZ) for up to 5 weeks, to induce myelin degeneration. Demyelination was monitored by histopathological analysis in different animal cohorts (7-10 animals/group) at weeks 3 and 5 of cuprizone diet (demyelination in A-C). One of the cuprizone diet cohorts was placed in a regular food diet (ND, normal diet) for additional 8 days to allow for spontaneous remyelination (CPZ week 5 + ND 8 days). Global lipidic staining with luxol fast blue (LFB, A) as well as immunodetection for myelin oligodendrocyte glycoprotein (MOG, B) and the mature oligodendrocyte marker GSTpi (C), were performed in coronal brain sections containing the corpus callosum. Area covered by LFB and MOG was used to determine the extent of de/remyelination while GSTpi allowed for accurate quantification of oligodendrocytes in the tissue sections. Data are shown as mean ± SEM of 1-2 sections per mouse. ***: p<0.05 versus the corresponding condition at week 3, according to one-way ANOVA. No statistical differences were found between conditions in each time point.

## Discussion

A matter of continuous debate in the myelin and oligodendrocyte biology field is whether the molecular and cellular mechanisms driving OPC maturation and myelin formation in early developmental stages are similar and follow the same patterns as the ones needed to make *de novo* myelin in homeostasis or lesion repair in the adult CNS. A great deal of current knowledge on the biological processes that modulate or influence myelination is the result of decades of studies in cell models that utilize OPCs isolated from postnatal rodent brains (Butt, Papanikolaou, & Rivera, 2019; Michalski, & Kothary, 2015). This has shed light on the intrinsic cellular changes taking place in the OPC journey to become myelinating cells during development, but at the same time it has effectively biased scientific assumptions for remyelination processes. A better understanding of the nature of OPC maturation in demyelinated lesions, considering the presence of active inflammation and non-cell autonomous mechanisms, is necessary for advancing effective therapies aimed to repair damaged myelin in diseases such as MS.

GPR17 is expressed at high levels in early stages of OPC maturation and it has been described to enhance developmental maturation and remyelination after genetic deletion or upon modulation of its function with surrogate ligands (Chen et al., 2009; Hennen et al., 2013). Several reports have therefore characterized this GPCR as a fundamental regulator of OPC maturation and proposed it as a druggable target for remyelination therapies (Hennen et al., 2013; Merten et al., 2018; Ou et al., 2016). We have not been able to corroborate some of those findings. First, upon performing an extensive analysis of GPR17-related gene signature by next generation RNA sequencing, we observed a marginal increase of myelin proteins when comparing GPR17^-/-^ to wild type cells. Interestingly, in this analysis prominent genes being upregulated in the absence of the receptor have, to our knowledge, no known functional relation to those driving OPC maturation and myelin formation. Confirming these observations from the gene expression perspective, we experimentally determined that GPR17 absence marginally modulates OPC maturation in postnatal cultures.

GPR17 is an orphan receptor reported to be antagonized by cysteinyl leukotriene receptor antagonists (e.g. pranlukast, montelukast) in the presence of surrogate agonists such as MDL29951, identified in small molecule screenings (Hennen et al., 2013). These compounds had the ability to modulate GPR17 function in cellular expression systems. We and others have observed receptor internalization upon agonistic binding in OPC cultures, but their effect on maturation was minor in terms of the transition from bipolar OPCs into MBP^+^ cells and beyond into actively myelinating cells. Furthermore, the marginal effect of GPR17 activity modulation on OPCs was similar in cells at different stages of maturation. This is important as it has been reported that OPCs in pre-myelinating stages respond differently to GPR17 modulation than earlier progenitors and that prolonged GPR17 activation leads to receptor internalization (Fratangeli et al., 2013; Parravicini et al., 2020).

Although we have confirmed an elevated expression of GPR17 during early stages of remyelination in experimentally induced demyelination (Nyamoya et al., 2019), our results do not support previous reports where antagonizing the receptor with pranlukast in a lysolecithin-demyelinated corpus callosum paradigm, significantly enhanced myelin recovery (Ou et al., 2016). In our hands, pranlukast showed no brain penetration in mouse pharmacokinetic studies (data not shown) indicating that its beneficial effects on myelin recovery were at best the result of indirect modulation of alternative signaling pathways in the periphery. Similarly, pharmacological modulation of GPR17 has been reported to marginally ameliorate disease scores in the EAE (experimental autoimmune encephalomyelitis) mouse model of MS (Parravicini et al., 2020). This data likely reflects modulation of the autoimmune response rather than a direct effect on myelin recovery. Furthermore, using genetically modified mice and two different de/remyelination animal models in the absence of peripheral inflammation, we were not able to determine any differences in the ratio of myelin recovery when comparing between wild type and GPR17^-/-^.

We have generated pharmacological data on GPR17 for several compounds reported to modulate OPC maturation as well as novel small molecule families identified from a Sanofi library wide screen. However, only minor functional OPC maturation effects were detected in response to the most potent compounds, which falls in line with the fact that the level of enhanced maturation shown in previous publications by modulators of the receptor is marginal. Additionally, though beyond the scope of our studies, there is also a high probability of off target effects triggered by these chemicals which could be having a direct impact on OPC maturation independently of GPR17 function.

One aspect of GPR17 biology to keep in consideration is the beneficial modulation of survival described not only for cells of the oligodendrocyte lineage exposed to LPC (Lu et al., 2018), but also for neuronal cells in rejuvenation studies (Marschallinger et al., 2015). This may have cellular implications extending from OPC survival to myelin preservation during injury or disease, and generate contradictory experimental outcomes easily interpreted as enhanced myelination/remyelination. A compelling demonstration of GPR17 function in OPC survival was observed in a mouse model of amyotrophic lateral sclerosis (ALS) where early upregulation of GPR17 appears to promote progenitor regeneration. However, myelination is impaired when GPR17 expression is maintained in mature cells for long periods (Bonfanti et al., 2020).

In the search for effective therapies for myelin repair in demyelinating diseases, it is of paramount importance to identify and fully validate the right biology on the right molecular target, using effective pharmacological tools, to then enable the development of drugs with adequate safety and efficacy. We believe GPR17 has a function in early OPC biology that may be related to cell survival and, to a lesser extent, to a developmental, dispensable enabler of maturation. It appears however, to lack the strength to regulate on its own the transition from OPCs into myelinating oligodendrocytes.

## Acknowledgements

The authors would like to thank Amy Mahan (Sanofi) for processing and analyzing real-time PCR using MS brain tissue samples, Theresa Kuntzweiler (Sanofi) for advice and data processing and Cynthia Pryce (Sanofi) for assistance with image analysis.

## Data Availability Statement

The data that support the findings of this study are available from the corresponding author upon reasonable request.

## References

1. Acs, P., Selak, M. A., Komoly, S., & Kalman, B. (2013). Distribution of oligodendrocyte loss and mitochondrial toxicity in the cuprizone-induced experimental demyelination model. J Neuroimmunol, 262(1-2), 128–131. doi:10.1016/j.jneuroim.2013.06.012

2. Anastasiadou, S., Liebenehm, S., Sinske, D., Meyer zu Reckendorf, C., Moepps, B., Nordheim, A., & Knoll, B. (2015). Neuronal expression of the transcription factor serum response factor modulates myelination in a mouse multiple sclerosis model. Glia, 63(6), 958–976. doi:10.1002/glia.22794

3. Barres, B. A., Lazar, M. A., & Raff, M. C. (1994). A novel role for thyroid hormone, glucocorticoids and retinoic acid in timing oligodendrocyte development. Development, 120(5), 1097–1108. Retrieved from https://www.ncbi.nlm.nih.gov/pubmed/8026323

4. Baumann, N., & Pham-Dinh, D. (2001). Biology of oligodendrocyte and myelin in the mammalian central nervous system. Physiol Rev, 81(2), 871–927. doi:10.1152/physrev.2001.81.2.871

5. Bläsius, R., Weber, R. G., Lichter, P., & Ogilvie, A. (1998). A Novel Orphan G Protein-Coupled Receptor Primarily Expressed in the Brain Is Localized on Human Chromosomal Band 2q21. 70(4), 1357–1365. doi:10.1046/j.1471-4159.1998.70041357.x

6. Boda, E., Vigano, F., Rosa, P., Fumagalli, M., Labat-Gest, V., Tempia, F., … Buffo, A. (2011). The GPR17 receptor in NG2 expressing cells: focus on in vivo cell maturation and participation in acute trauma and chronic damage. Glia, 59(12), 1958–1973. doi:10.1002/glia.21237

7. Bonfanti, E., Bonifacino, T., Raffaele, S., Milanese, M., Morgante, E., Bonanno, G., … Fumagalli, M. (2020). Abnormal Upregulation of GPR17 Receptor Contributes to Oligodendrocyte Dysfunction in SOD1 G93A Mice. Int J Mol Sci, 21(7). doi:10.3390/ijms21072395

8. Boyd, A., Zhang, H., & Williams, A. (2013). Insufficient OPC migration into demyelinated lesions is a cause of poor remyelination in MS and mouse models. Acta Neuropathol, 125(6), 841–859. doi:10.1007/s00401-013-1112-y

9. Butt, A. M., Papanikolaou, M., & Rivera, A. (2019). Physiology of Oligodendroglia. Adv Exp Med Biol, 1175, 117–128. doi:10.1007/978-981-13-9913-8_5

10. Capra, V., Thompson, M. D., Sala, A., Cole, D. E., Folco, G., & Rovati, G. E. (2007). Cysteinyl-leukotrienes and their receptors in asthma and other inflammatory diseases: critical update and emerging trends. Med Res Rev, 27(4), 469–527. doi:10.1002/med.20071

11. Ceruti, S., Vigano, F., Boda, E., Ferrario, S., Magni, G., Boccazzi, M., … Abbracchio, M. P. (2011). Expression of the new P2Y-like receptor GPR17 during oligodendrocyte precursor cell maturation regulates sensitivity to ATP-induced death. Glia, 59(3), 363–378. doi:10.1002/glia.21107

12. Chen, Y., Wu, H., Wang, S., Koito, H., Li, J., Ye, F., … Lu, Q. R. (2009). The oligodendrocyte-specific G protein-coupled receptor GPR17 is a cell-intrinsic timer of myelination. Nat Neurosci, 12(11), 1398–1406. doi:10.1038/nn.2410

13. Ciana, P., Fumagalli, M., Trincavelli, M. L., Verderio, C., Rosa, P., Lecca, D., … Abbracchio, M. P. (2006). The orphan receptor GPR17 identified as a new dual uracil nucleotides/cysteinyl-leukotrienes receptor. EMBO J, 25(19), 4615–4627. doi:10.1038/sj.emboj.7601341

14. Coclitu, C., Constantinescu, C. S., & Tanasescu, R. (2016). The future of multiple sclerosis treatments. Expert Rev Neurother, 16(12), 1341–1356. doi:10.1080/14737175.2016.1243056

15. Dugas, J. C., & Emery, B. (2013). Purification of oligodendrocyte precursor cells from rat cortices by immunopanning. Cold Spring Harb Protoc, 2013(8), 745–758. doi:10.1101/pdb.prot070862

16. Dziedzic, A., Miller, E., Saluk-Bijak, J., & Bijak, M. (2020). The GPR17 Receptor-A Promising Goal for Therapy and a Potential Marker of the Neurodegenerative Process in Multiple Sclerosis. Int J Mol Sci, 21(5). doi:10.3390/ijms21051852

17. Fancy, S. P., Kotter, M. R., Harrington, E. P., Huang, J. K., Zhao, C., Rowitch, D. H., & Franklin, R. J. (2010). Overcoming remyelination failure in multiple sclerosis and other myelin disorders. Exp Neurol, 225(1), 18–23. doi:10.1016/j.expneurol.2009.12.020

18. Ferguson, S. S. (2001). Evolving concepts in G protein-coupled receptor endocytosis: the role in receptor desensitization and signaling. Pharmacol Rev, 53(1), 1–24. Retrieved from https://www.ncbi.nlm.nih.gov/pubmed/11171937

19. Franklin, R. J., & Ffrench-Constant, C. (2008). Remyelination in the CNS: from biology to therapy. Nat Rev Neurosci, 9(11), 839–855. doi:10.1038/nrn2480

20. Fratangeli, A., Parmigiani, E., Fumagalli, M., Lecca, D., Benfante, R., Passafaro, M., … Rosa, P. (2013). The regulated expression, intracellular trafficking, and membrane recycling of the P2Y-like receptor GPR17 in Oli-neu oligodendroglial cells. J Biol Chem, 288(7), 5241–5256. doi:10.1074/jbc.M112.404996

21. Fumagalli, M., Daniele, S., Lecca, D., Lee, P. R., Parravicini, C., Fields, R. D., … Abbracchio, M. P. (2011). Phenotypic changes, signaling pathway, and functional correlates of GPR17-expressing neural precursor cells during oligodendrocyte differentiation. J Biol Chem, 286(12), 10593–10604. doi:10.1074/jbc.M110.162867

22. Goldschmidt, T., Antel, J., König, F. B., Brück, W., & Kuhlmann, T. (2009). Remyelination capacity of the MS brain decreases with disease chronicity. 72(22), 1914–1921. doi:10.1212/WNL.0b013e3181a8260a %J Neurology

23. Green, A. J., Gelfand, J. M., Cree, B. A., Bevan, C., Boscardin, W. J., Mei, F., … Chan, J. R. (2017). Clemastine fumarate as a remyelinating therapy for multiple sclerosis (ReBUILD): a randomised, controlled, double-blind, crossover trial. Lancet, 390(10111), 2481–2489. doi:10.1016/S0140-6736(17)32346-2

24. Hall, S. M. (1972). The effect of injections of lysophosphatidyl choline into white matter of the adult mouse spinal cord. J Cell Sci, 10(2), 535–546.

25. Harlow, D. E., Honce, J. M., & Miravalle, A. A. (2015). Remyelination Therapy in Multiple Sclerosis. Front Neurol, 6, 257. doi:10.3389/fneur.2015.00257

26. Hennen, S., Wang, H., Peters, L., Merten, N., Simon, K., Spinrath, A., … Kostenis, E. (2013). Decoding signaling and function of the orphan G protein-coupled receptor GPR17 with a small-molecule agonist. Sci Signal, 6(298), ra93. doi:10.1126/scisignal.2004350

27. Hiremath, M. M., Saito, Y., Knapp, G. W., Ting, J. P., Suzuki, K., & Matsushima, G. K. (1998). Microglial/macrophage accumulation during cuprizone-induced demyelination in C57BL/6 mice. J Neuroimmunol, 92(1-2), 38–49. doi:10.1016/s0165-5728(98)00168-4

28. Hooijmans, C. R., Hlavica, M., Schuler, F. A. F., Good, N., Good, A., Baumgartner, L., … Ineichen, B. V. (2019). Remyelination promoting therapies in multiple sclerosis animal models: a systematic review and meta-analysis. Sci Rep, 9(1), 822. doi:10.1038/s41598-018-35734-4

29. Jeffery, N. D., & Blakemore, W. F. (1995). Remyelination of mouse spinal cord axons demyelinated by local injection of lysolecithin. J Neurocytol, 24(10), 775–781. doi:10.1007/bf01191213

30. Jurevics, H., Largent, C., Hostettler, J., Sammond, D. W., Matsushima, G. K., Kleindienst, A., … Morell, P. (2002). Alterations in metabolism and gene expression in brain regions during cuprizone-induced demyelination and remyelination. J Neurochem, 82(1), 126–136. doi:10.1046/j.1471-4159.2002.00954.x

31. Kolahdouzan, M., Futhey, N. C., Kieran, N. W., & Healy, L. M. (2019). Novel Molecular Leads for the Prevention of Damage and the Promotion of Repair in Neuroimmunological Disease. 10(1657). doi:10.3389/fimmu.2019.01657

32. Kremer, D., Gottle, P., Flores-Rivera, J., Hartung, H. P., & Kury, P. (2019). Remyelination in multiple sclerosis: from concept to clinical trials. Curr Opin Neurol, 32(3), 378–384. doi:10.1097/WCO.0000000000000692

33. Kuhlmann, T., Miron, V., Cuo, Q., Wegner, C., Antel, J., & Brück, W. (2008). Differentiation block of oligodendroglial progenitor cells as a cause for remyelination failure in chronic multiple sclerosis. Brain, 131(7), 1749–1758. doi:10.1093/brain/awn096 %J Brain

34. Lassmann, H., Bruck, W., & Lucchinetti, C. F. (2007). The immunopathology of multiple sclerosis: an overview. Brain Pathol, 17(2), 210–218. doi:10.1111/j.1750-3639.2007.00064.x

35. Lecca, D., Trincavelli, M. L., Gelosa, P., Sironi, L., Ciana, P., Fumagalli, M., … Abbracchio, M. P. (2008). The recently identified P2Y-like receptor GPR17 is a sensor of brain damage and a new target for brain repair. PLoS One, 3(10), e3579. doi:10.1371/journal.pone.0003579

36. Lu, C., Dong, L., Zhou, H., Li, Q., Huang, G., Bai, S. J., & Liao, L. (2018). G-Protein-Coupled Receptor Gpr17 Regulates Oligodendrocyte Differentiation in Response to Lysolecithin-Induced Demyelination. Sci Rep, 8(1), 4502. doi:10.1038/s41598-018-22452-0

37. Marchese, A., Paing, M. M., Temple, B. R., & Trejo, J. (2008). G protein-coupled receptor sorting to endosomes and lysosomes. Annu Rev Pharmacol Toxicol, 48, 601–629. doi:10.1146/annurev.pharmtox.48.113006.094646

38. Marschallinger, J., Schaffner, I., Klein, B., Gelfert, R., Rivera, F. J., Illes, S., … Aigner, L. (2015). Structural and functional rejuvenation of the aged brain by an approved anti-asthmatic drug. Nat Commun, 6, 8466. doi:10.1038/ncomms9466

39. Marucci, G., Dal Ben, D., Lambertucci, C., Santinelli, C., Spinaci, A., Thomas, A., … Buccioni, M. (2016). The G Protein-Coupled Receptor GPR17: Overview and Update. ChemMedChem, 11(23), 2567–2574. doi:10.1002/cmdc.201600453

40. Mastaitis, J., Min, S., Elvert, R., Kannt, A., Xin, Y., Ochoa, F., … Gromada, J. (2015). GPR17 gene disruption does not alter food intake or glucose homeostasis in mice. Proc Natl Acad Sci U S A, 112(6), 1845–1849. doi:10.1073/pnas.1424968112

41. Matsushima, G. K., & Morell, P. (2001). The neurotoxicant, cuprizone, as a model to study demyelination and remyelination in the central nervous system. Brain Pathol, 11(1), 107–116. doi:10.1111/j.1750-3639.2001.tb00385.x

42. Mei, F., Fancy, S. P. J., Shen, Y. A., Niu, J., Zhao, C., Presley, B., … Chan, J. R. (2014). Micropillar arrays as a high-throughput screening platform for therapeutics in multiple sclerosis. Nat Med, 20(8), 954–960. doi:10.1038/nm.3618

43. Merten, N., Fischer, J., Simon, K., Zhang, L., Schroder, R., Peters, L., … Kostenis, E. (2018). Repurposing HAMI3379 to Block GPR17 and Promote Rodent and Human Oligodendrocyte Differentiation. Cell Chem Biol, 25(6), 775–786.e775. doi:10.1016/j.chembiol.2018.03.012

44. Michalski, J.-P., & Kothary, R. (2015). Oligodendrocytes in a Nutshell. Frontiers in cellular neuroscience, 9, 340–340. doi:10.3389/fncel.2015.00340

45. Miron, V. E., Kuhlmann, T., & Antel, J. P. (2011). Cells of the oligodendroglial lineage, myelination, and remyelination. Biochim Biophys Acta, 1812(2), 184–193. doi:10.1016/j.bbadis.2010.09.010

46. Montalban, X., Hauser, S. L., Kappos, L., Arnold, D. L., Bar-Or, A., Comi, G., … Wolinsky, J. S. (2017). Ocrelizumab versus Placebo in Primary Progressive Multiple Sclerosis. N Engl J Med, 376(3), 209–220. doi:10.1056/NEJMoa1606468

47. Motavaf, M., Sadeghizadeh, M., & Javan, M. (2017). Attempts to Overcome Remyelination Failure: Toward Opening New Therapeutic Avenues for Multiple Sclerosis. Cell Mol Neurobiol, 37(8), 1335–1348. doi:10.1007/s10571-017-0472-6

48. Najm, F. J., Madhavan, M., Zaremba, A., Shick, E., Karl, R. T., Factor, D. C., … Tesar, P. J. (2015). Drug-based modulation of endogenous stem cells promotes functional remyelination in vivo. Nature, 522(7555), 216–220. doi:10.1038/nature14335

49. Nyamoya, S., Leopold, P., Becker, B., Beyer, C., Hustadt, F., Schmitz, C., … Kipp, M. (2019). G-Protein-Coupled Receptor Gpr17 Expression in Two Multiple Sclerosis Remyelination Models. Mol Neurobiol, 56(2), 1109–1123. doi:10.1007/s12035-018-1146-1

50. Olsen, J. A., & Akirav, E. M. (2015). Remyelination in multiple sclerosis: cellular mechanisms and novel therapeutic approaches. J Neurosci Res, 93(5), 687–696. doi:10.1002/jnr.23493

51. O’Meara, R. W., Ryan, S. D., Colognato, H., & Kothary, R. (2011). Derivation of enriched oligodendrocyte cultures and oligodendrocyte/neuron myelinating co-cultures from post-natal murine tissues. J Vis Exp(54). doi:10.3791/3324

52. Ou, Z., Sun, Y., Lin, L., You, N., Liu, X., Li, H., … Chen, Y. (2016). Olig2-Targeted G-Protein-Coupled Receptor Gpr17 Regulates Oligodendrocyte Survival in Response to Lysolecithin-Induced Demyelination. J Neurosci, 36(41), 10560–10573. doi:10.1523/JNEUROSCI.0898-16.2016

53. Parravicini, C., Lecca, D., Marangon, D., Coppolino, G. T., Daniele, S., Bonfanti, E., … Eberini, I. (2020). Development of the first in vivo GPR17 ligand through an iterative drug discovery pipeline: A novel disease-modifying strategy for multiple sclerosis. PLoS One, 15(4), e0231483. doi:10.1371/journal.pone.0231483

54. Praet, J., Guglielmetti, C., Berneman, Z., Van der Linden, A., & Ponsaerts, P. (2014). Cellular and molecular neuropathology of the cuprizone mouse model: clinical relevance for multiple sclerosis. Neurosci Biobehav Rev, 47, 485–505. doi:10.1016/j.neubiorev.2014.10.004

55. Rajagopal, S., & Shenoy, S. K. (2018). GPCR desensitization: Acute and prolonged phases. Cell Signal, 41, 9–16. doi:10.1016/j.cellsig.2017.01.024

56. Raport, C. J., Schweickart, V. L., Chantry, D., Eddy Jr., R. L., Shows, T. B., Godiska, R., & Gray, P. W. (1996). New members of the chemokine receptor gene family. 59(1), 18–23. doi:10.1002/jlb.59.1.18

57. Rommer, P. S., Milo, R., Han, M. H., Satyanarayan, S., Sellner, J., Hauer, L., … Stuve, O. (2019). Immunological Aspects of Approved MS Therapeutics. 10(1564). doi:10.3389/fimmu.2019.01564

58. Schwartzbach, C. J., Grove, R. A., Brown, R., Tompson, D., Then Bergh, F., & Arnold, D. L. (2017). Lesion remyelinating activity of GSK239512 versus placebo in patients with relapsing-remitting multiple sclerosis: a randomised, single-blind, phase II study. J Neurol, 264(2), 304–315. doi:10.1007/s00415-016-8341-7

59. Simon, K., Hennen, S., Merten, N., Blattermann, S., Gillard, M., Kostenis, E., & Gomeza, J. (2016). The Orphan G Protein-coupled Receptor GPR17 Negatively Regulates Oligodendrocyte Differentiation via Galphai/o and Its Downstream Effector Molecules. J Biol Chem, 291(2), 705–718. doi:10.1074/jbc.M115.683953

60. Stangel, M., Kuhlmann, T., Matthews, P. M., & Kilpatrick, T. J. (2017). Achievements and obstacles of remyelinating therapies in multiple sclerosis. Nat Rev Neurol, 13(12), 742–754. doi:10.1038/nrneurol.2017.139

61. Trapp, B. D., Peterson, J., Ransohoff, R. M., Rudick, R., Mörk, S., & Bö, L. (1998). Axonal Transection in the Lesions of Multiple Sclerosis. 338(5), 278–285. doi:10.1056/nejm199801293380502

62. Zhang, Y., Zhang, H., Wang, L., Jiang, W., Xu, H., Xiao, L., … Li, X. M. (2012). Quetiapine enhances oligodendrocyte regeneration and myelin repair after cuprizone-induced demyelination. Schizophr Res, 138(1), 8–17. doi:10.1016/j.schres.2012.04.006

